# Beyond Redfield: Thermodynamic Bounds and Non-Perturbative Quantum Dynamics in Tubulin Networks

**DOI:** 10.64898/2026.05.10.724047

**Authors:** Facundo Firmenich, Pau Firmenich, León Firmenich

**Affiliations:** Centro de Estudios del Sur (CEDESUR), Spain and Argentina. For this project, academically affiliated with Universidad Nacional Arturo Jauretche (UNAJ), Av. Calchaquí 6200, Florencio Varela, Provincia de Buenos Aires, 1888, Argentina; Universitat de Barcelona (UB), Gran Via de les Corts Catalanes, 585, 08007 Barcelona, Spain

**Keywords:** biological physics, quantum coherence, microtubules, tubulin, open quantum systems, HEOM, Redfield theory, Fröhlich condensation, QED cavity, subradiance, quantum discord, neural synchronization, quantum biology

## Abstract

Quantum effects in biology are unavoidable at the molecular scale; the unresolved question is whether they can remain functionally relevant across the timescale gap between femtosecond molecular dynamics and microsecond-to-millisecond biological function. Here we formalize this mismatch as an *equilibrium-to-functionality gap* and use tubulin as a stringent open-system test case.

We combine secular Lindblad, Redfield, and hierarchical equations of motion (HEOM) treatments to quantify decoherence, non-perturbative relaxation, and the physical amplification required for functional relevance. Equilibrium dephasing yields a conservative 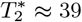 fs at 310 K, with a generic protein-bath baseline of ≈ 13 fs. A completed 30 ps HEOM trajectory for the full 1JFF tryptophan network shows distributed non-Markovian relaxation, with terminal purity Pur = 0.210 and stretched-exponential exponent *β*_KWW_ ≈ 0.44, confirming that Redfield is useful as a short-time perturbative comparator but not quantitatively interchangeable with HEOM in this intermediate-coupling regime.

We introduce a coherence-utility criterion 𝒰 = 𝒦*τ*_coh_*/τ*_func_, separating required amplification from empirically bounded gain. A thermodynamic uncertainty relation closure shows that neural-scale cascade amplification would require *P*_min_ ∼ 10^−7^ W, about five orders of magnitude above the local microtubule GTP budget. Fröhlich pumping is found to be linewidth-gated rather than generically micron-scale; ordered-water cavity QED and geometric subradiance remain experimentally testable but severely constrained candidates.

The result is not a model of consciousness, but a reproducible physical benchmark framework for evaluating biological quantum-coherence claims under explicit open-system, energetic, and experimental constraints. Six falsifiable experimental programmes are prioritized, and the full computational framework is released with a validation ledger, cryptographic audit trail, and living supplementary material.

Graphical abstract.
Equilibrium tubulin coherence lies in the femtosecond regime, while functional neural timescales lie in the millisecond regime. Fröhlich pumping, QED-cavity protection, and geometric subradiance remain experimentally discriminable non-equilibrium candidates requiring independently bounded amplification.

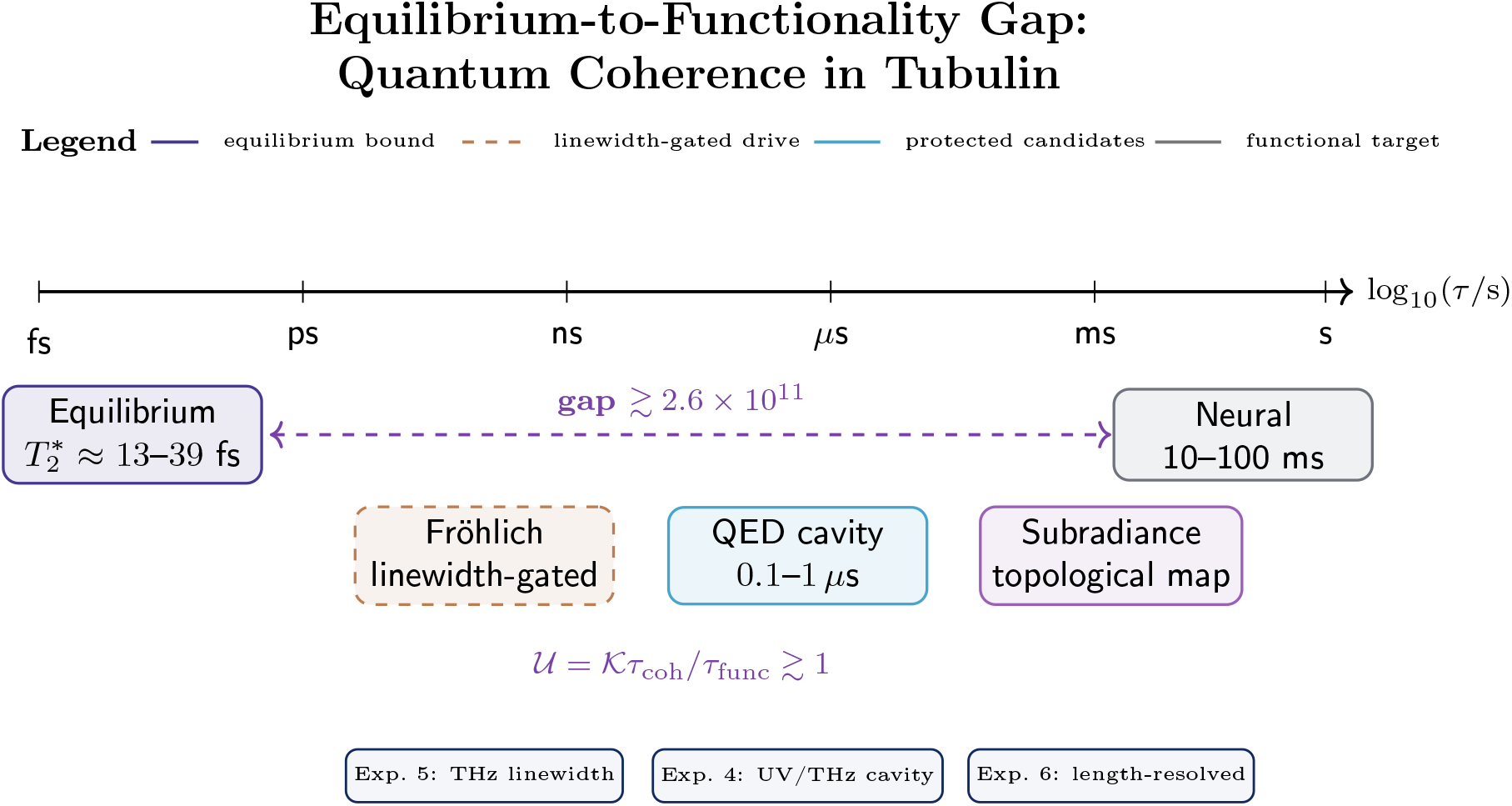

**Funding:** This research received no specific grant from any funding agency in the public, commercial, or not-for-profit sectors.

**Versioned computational scope of this release:** This manuscript reports the theoretical framework, calibrated equilibrium baseline, Redfield/HEOM validation ledger, stratified Bayesian evidence synthesis, classical comparators, and falsifiable experimental design. The release-specific reproduction audit, including the current validation-check total and the SHA-256 fingerprints of the binary production artefacts (`.npz`, `.pkl`), is documented in `LIVING_SI.md` and `outputs_data/raw_json/structur al/validation_report.json`. A completed 30 ps HEOM production trajectory has been validated on constrained hardware; the master dataset contains the full 8-site population trajectory. A summary of those results is provided in §<u>2.2.5</u>. All claims made below are restricted to the numerical and theoretical evidence reported in this manuscript and its associated repository artefacts. The public repository ships the calibrated phenomenological baseline for accessibility; the HEOM production artefacts serve as the non-perturbative validation benchmark. All source figure outputs associated with this release are maintained in the public repository under `outputs_data/figures_final/`.

## 1 Introduction: biological quantum relevance as an equilibrium-to-functionality problem

The central question addressed here is not whether quantum mechanics is present in biomolecules—it necessarily is—but whether molecular quantum dynamics can remain operationally relevant across the enormous timescale separation between femtosecond/picosecond molecular motion and microsecond-to-millisecond biological function. We treat this separation as an *equilibrium-to-functionality gap*. The gap is especially severe for tubulin, which is used here not because it is assumed to be privileged, but because it is a stringent and historically important stress test: structurally ordered, optically rich, biologically central, and embedded in a warm, wet, strongly fluctuating cellular environment.

Our objective is deliberately conservative. We do not assume that tubulin quantum coherence contributes to neural computation, nor do we attempt to explain consciousness. Indeed, the coherence utility framework 𝒰 introduced here is entirely independent of any specific consciousness model, including Orch OR [69, 70]; it addresses the narrower, operationally precise question of whether molecular quantum dynamics can produce physically distinguishable signatures on functionally relevant timescales. Instead, we ask what physical conditions would be required for molecular quantum dynamics in tubulin to become functionally consequential, and which experiments could rule in or rule out the relevant non-equilibrium model classes.

### 1.1 Quantum coherence in biology: revised understanding (2017–2025)

Quantum mechanics was traditionally considered irrelevant for biology beyond bond formation. This shifted with demonstrations of quantum effects in three established systems plus one emerging one. **Photosynthetic light harvesting (FMO complex)**. Initial reports [1, 2] of *τ* ∼ 300–600 fs electronic coherence at 277 K have been substantially revised. Thyrhaug et al. [4] demonstrated using polarization-controlled 2D spectroscopy that long-lived quantum beats are exclusively vibrational in origin, whereas electronic coherences dephase within 240 fs even at 77 K. At physiological temperature, Duan et al. [3] found electronic coherence *τ*_elec_ *<* 60 fs at 295 K. Computational studies [6, 7, 8] showed vibrational nuclear motion produces spectral signatures previously attributed to electronic coherence. Thyrhaug et al. [4, 5] confirmed electronic coherences are ultrafast (∼ 60 fs) at room temperature for high-energy excitons, extending to ∼ 500 fs only at cryogenic temperatures. **Functional implications**. Despite ultrafast electronic decoherence, energy transfer in FMO occurs over ∼ 1 ps timescales, suggesting vibrational coherences and environment-assisted quantum transport (ENAQT) [10] contribute to efficiency even without persistent electronic coherence [11]. The quantum advantage appears to lie in vibronic coupling [12, 9] and correlated environmental fluctuations rather than long-lived electronic superpositions. **Avian magnetoreception**. Radical pair spin coherence in cryptochrome proteins exhibits *τ* ∼ 1–10 *µ*s at 300 K with experimental validation via magnetic isotope effects [13, 14, 15]. **Enzymatic tunneling**. Kinetic isotope effects (KIE ∼ 3–80) in hydrogen transfer enzymes indicate quantum tunneling contributions to catalysis [16, 17]. **Microtubule tryptophan networks**. Babcock et al. [18] experimentally demonstrated UV superradiance in mega-networks (*>* 10^5^ tryptophan dipoles) of microtubule architectures at room temperature, building on theoretical predictions of superradiant transport in tubulin [19]. Superradiant states (*τ* ∼ 100 fs) coexist with assigned long-lived subradiant components (*τ* ∼ 10 s), providing direct experimental evidence consistent with collective quantum-optical effects in MT architectures.

### 1.2 The microtubule hypothesis: QED cavity and non-equilibrium models

Microtubules are hollow cylinders (outer diameter 25 nm, inner lumen 15 nm, length 1–50 *µ*m) assembled from *αβ*-tubulin dimers in 13 protofilaments. Each neuron contains ∼ 10^7^ dimers. **QED cavity model**. Mavromatos, Mershin, and Nanopoulos [20, 21, 22] developed a model treating MT interiors (containing ordered water [23, 24]) as high-Q electromagnetic cavities. Strong dipole-dipole coupling between tubulin dimers (*µ* ∼ 1700 Debye [25]) and ordered water dipoles in a ∼ 1 nm hydrophobic wall layer provides environmental shielding. Decoherence arises primarily from leakage of water dipole quanta, yielding:

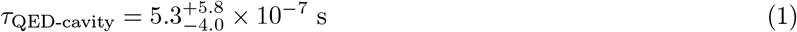

with scaling *τ* ∝ *ε*^2^, where *ε* = 80 ± 20 is the ordered-water dielectric constant (Monte Carlo propagation, 95% CI). **Non-equilibrium (Fröhlich) models**. Fröhlich [26, 27] demonstrated that dipole systems under continuous metabolic pumping can form Bose-like condensates at high temperature, explicitly violating equilibrium thermodynamics. For MT dimers continuously supplied with GTP hydrolysis energy (Δ*E* ∼ 0.42 eV ≈ 16*k*_B_*T* at 310 K), coherence of a collective mode at frequency *ω*_*F*_ could be sustained with:

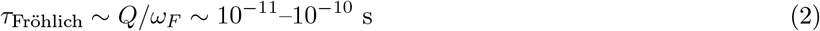

where *Q* is the quality factor of the condensed mode. **Controversy**. Reimers et al. [28] analyzed one-dimensional polymers and concluded that Fröhlich condensates are inaccessible in a biological environment. However, Pokorný et al. [29] noted that this analysis uses linearized equations neglecting non-linear elastic-electric coupling central to Fröhlich’s original framework, and models 1D chains rather than cylindrical MT geometry with ordered water [20, 30]. The discrepancy between these analyses defines the central unresolved theoretical question, which we recast quantitatively in §4.1 by showing that micron-scale gating is linewidth-conditional rather than a direct consequence of the THz carrier frequency. **Recent experimental support**. Khan et al. [32] showed MT-stabilizing drugs delayed anesthetic-induced unconsciousness in rats (Cohen’s *d* = 1.9). Kalra et al. [33] demonstrated anesthetics dampen quantum optical effects in MTs. Wiest [34] reviewed convergent evidence that volatile anesthetics act on MT quantum states. Saxena et al. [35] and Singh et al. [36] observed MT resonances spanning multiple neurons.

### 1.3 Equilibrium vs non-equilibrium: the critical distinction

Tegmark’s critique [37] assumed thermal equilibrium and large spatial superpositions. Hagan et al. [38] corrected the spatial assumption. The fundamental issue remains: equilibrium assumptions are inappropriate for living systems continuously dissipating metabolic energy. Our approach: §§2–3 establish rigorous equilibrium bounds (*τ* ∼ ps). §3.4 introduces a novel coherence utility framework quantifying the amplification gap. §4 analyzes non-equilibrium mechanisms, explicitly auditing finite-size Fröhlich gating and retaining cavity and subradiance only as conditionally constrained, falsifiable candidates. §5 provides a quantitative classical comparison with information-theoretic distinguishability criteria. §6 proposes six falsifiable experiments, hierarchized by discriminative power, with a four-tier validation architecture.

### 1.4 The equilibrium-to-functionality gap

Neural timescales (10–100 ms) vs molecular coherence yield two distinct scale gaps. First, carrying equilibrium fs coherence into a microsecond-scale cellular readout already requires 10^7^–10^8^ amplification. Second, bridging the same equilibrium baseline to neural-scale claims requires ∼ 10^11^–10^13^ amplification. If a non-equilibrium mechanism independently reaches the microsecond regime, the residual bridge to neural timing is still 10^4^–10^5^. For functional relevance, one of the following must hold: (1) extreme equilibrium protection yielding *τ* ∼ ms (unknown mechanism); cooperative amplification by factor ∼ *N* or 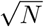; (3) non-equilibrium pumping sustaining coherence (Fröhlich); (4) cavity shielding by ordered water (Mavromatos et al.); (5) cascading quantum→classical amplification; (6) geometric decoherence-free subspaces (subradiant states); (7) classical mechanisms suffice (null hypothesis). We demonstrate equilibrium mechanisms (1–2) fail; Fröhlich (3) is linewidth-conditional and insufficient for neural-scale coherence under most conditions; non-equilibrium mechanisms (4, 6) are testable hypotheses.

### 1.5 Operational degree-of-freedom separation

A central separation used throughout this work is between conformational two-level-system (TLS) dynamics and tryptophan excitonic dynamics. The former defines the operational functional bound used in the coherence-utility criterion, whereas the latter provides a non-perturbative benchmark for structured open-system relaxation and a quantum-optical proxy for geometrically protected modes. Evidence for one degree of freedom is therefore not treated as evidence for the other unless an explicit transduction mechanism is specified.

### 1.6 Scope and contribution

**Objective 1 (§2):** Establish equilibrium decoherence constraints for tubulin using calibrated open-system models, spectral-density uncertainty, and Redfield/HEOM validation ledgers. **Objective 2 (§3):** Derive cooperative amplification bounds, introduce a quantum-coherence utility criterion distinguishing 𝒦_req_ from 𝒦_emp_, analyze quantum discord, and construct the Holstein vibronic analogue. **Objective 3 (§4):** Treat Fröhlich pumping as a linewidth-conditional finite-size mechanism, and ordered-water cavity QED and geometric subradiance as surviving non-equilibrium *model classes*. **Objective 4 (§5):** Compare candidate quantum mechanisms against known classical synchronization mechanisms and make the null hypothesis explicit. **Objective 5 (§6):** Provide six experimental programmes with explicit positive criteria, null criteria, confounds, and statistical decision layers, prioritized by discriminative power, with a four-tier validation architecture.

## 2 Equilibrium decoherence analysis

### 2.1 Two-level system model and the choice of open-system formalism

We model tubulin dimers as quantum two-state systems (“straight” |0⟩ vs “curved” |1⟩ conformations):

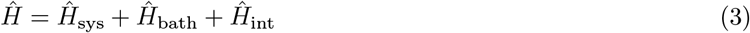

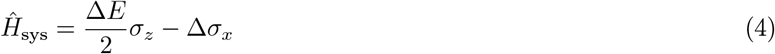

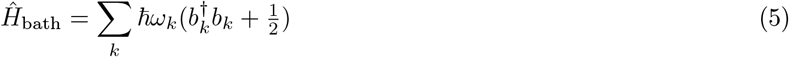

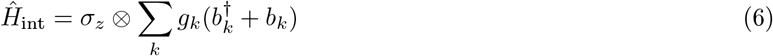

We adopt an Ohmic spectral density as the primary model:

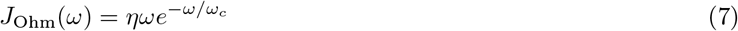

with dimensionless coupling *η* and cutoff *ω*_*c*_.

#### Choice of master equation

At *T* = 310 K with *η* ∼ 0.3 and *ω*_*c*_ ∼ 4.5 THz the relevant dimensionless parameters are *ħω*_*c*_*/k*_B_*T* ≈ 0.68 and 2*πηk*_B_*T/ħω*_*c*_ ≈ 1.4. The system therefore sits in the *intermediate-coupling / non-Markovian* regime [46, 44]. We use three levels of description:

1. *Secular Lindblad* — analytically tractable; used for closed-form scalings.
2. *Time-convolutionless Redfield* (second order) — retains non-secular coherence transfer; used for rate tables (Table 5).
3. *Hierarchical Equations of Motion* (HEOM) [47, 48] — numerically exact for Drude–Lorentz baths; used as benchmark.

**Table 1:**
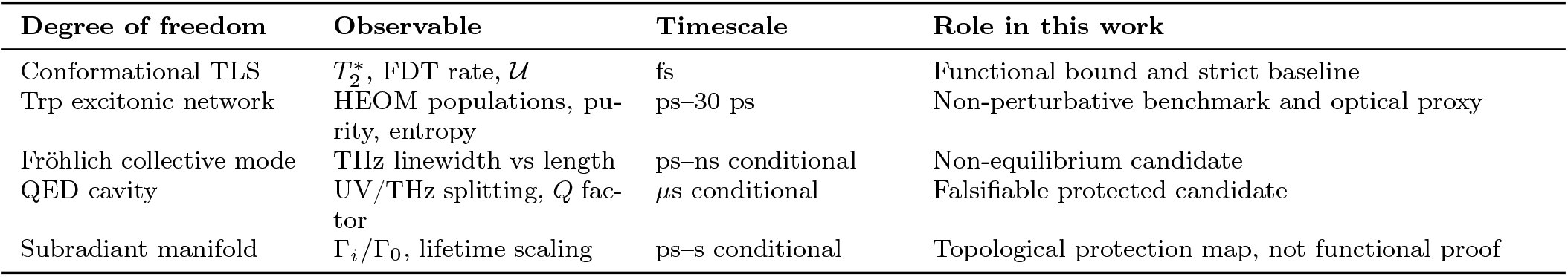
Operational roles of the degrees of freedom considered in this work.

**Table 2:** Reviewer quick map.

| Question | Where answered |
| --- | --- |
| What is the equilibrium coherence bound? | § 2 |
| Why does naive amplification fail? | § 3.1–3.4 |
| What is new beyond Redfield? | § 2.2.5, Appendix G, and Appendix N |
| What survives non-equilibrium scrutiny? | § 4 |
| What experiments decide the issue? | § 6 |
| What is reproduced computationally? | Appendix G and the Living SI |

**Table 3:** Production-scale HEOM benchmark and perturbative comparators for the full 1JFF tryptophan network.

| Method | Coherence scale | Diagnostic status |
| --- | --- | --- |
| Secular Lindblad | $\sim 2.3$ ps | Analytic Markovian reference; useful for closed-form scaling. |
| Redfield | $\sim 2.0$ ps | Perturbative short-time comparator; not quantitatively interchangeable with HEOM at intermediate coupling. |
| HEOM ($N_C = 7$, 500 fs benchmark) | $\sim 1.7$ ps | Non-perturbative short-window benchmark; confirms HEOM–Redfield model-level divergence. |
| HEOM ($N_C = 7$, 30 ps production) | — | Pur = 0.210, PR = 4.76, $S = 1.72 \pm 0.02$; $D_{\text{KL}} = 0.3809$ nats relative to Gibbs reference. |

**Table 4:** HEOM hierarchy convergence ledger for the 1JFF system.

| Comparison | $dPop$ | $dCoh$ | $dFrob$ |
| --- | --- | --- | --- |
| $N_C : 3 \rightarrow 4, N_k = 1$ | $5.51 \times 10^{-2}$ | $3.57 \times 10^{-3}$ | $9.97 \times 10^{-2}$ |
| $N_C : 4 \rightarrow 5, N_k = 1$ | $2.90 \times 10^{-2}$ | $2.87 \times 10^{-3}$ | $4.91 \times 10^{-2}$ |
| $N_C : 5 \rightarrow 6, N_k = 1$ | $1.29 \times 10^{-2}$ | $1.43 \times 10^{-3}$ | $2.24 \times 10^{-2}$ |
| $N_C : 6 \rightarrow 7, N_k = 1$ | $5.45 \times 10^{-3}$ | $9.10 \times 10^{-4}$ | $9.45 \times 10^{-3}$ |
| $N_C : 7 \rightarrow 8, N_k = 1$ | $2.09 \times 10^{-3}$ | $5.06 \times 10^{-4}$ | $3.76 \times 10^{-3}$ |
| $N_k : 1 \rightarrow 2, N_C = 5$ | $1.24 \times 10^{-4}$ | $3.69 \times 10^{-5}$ | $1.92 \times 10^{-4}$ |

**Table 5:** Decoherence rates in tubulin dimers at 310 K under equilibrium assumptions. Vibrational dephasing dominates. NYA = not yet attempted; AI = attempted, inconclusive; PS = partially supported; CR = consensus-robust.

| Source | Mechanism | Unprot. | Prot. | Protection | Scale | Experimental status |
| --- | --- | --- | --- | --- | --- | --- |
| Water | Collisional | $10^{13}$ | $10^{10}$ | Cavity, H<sub>2</sub>O order | Ext. | PS [24] |
| Ions | Electrostatic | $10^{10}$ | $10^9$ | Debye screening | Ext. | CR [43] |
| **Vibrations** | **Phonon coupling** | **$10^{13}$** | **$2.6 \times 10^{13}$** | **Rigidity, limited** | **Int.** | **NYA; direct tubulin 2D-IR proposed in §6.3** |
| Radiation | Spontaneous | $10^6$ | $10^6$ | None | Negl. | CR; standard QED |
| **Total** | — | $\sim 10^{13}$ | $\sim 2.6 \times 10^{13}$ | — | Vib-lim. | — |

**Table 6:** Subradiance threshold regimes under coherence-utility constraints.

| Regime | $\tau_A$ | $p$ required ($\mathcal{U} \geq 1$) | Status |
| --- | --- | --- | --- |
| Disorder-dominated | $\sim 10$ ps | $p > 0.25$ (not viable under bounds) | Ineffective |
| Partial geometric protection | $\sim 1$ μs | $p > 2.5 \times 10^{-3}$ | Marginal |
| Babcock-like | $\sim 10$ s | $p > 2.5 \times 10^{-9}$ | Broadly accessible |

**Table 7:**
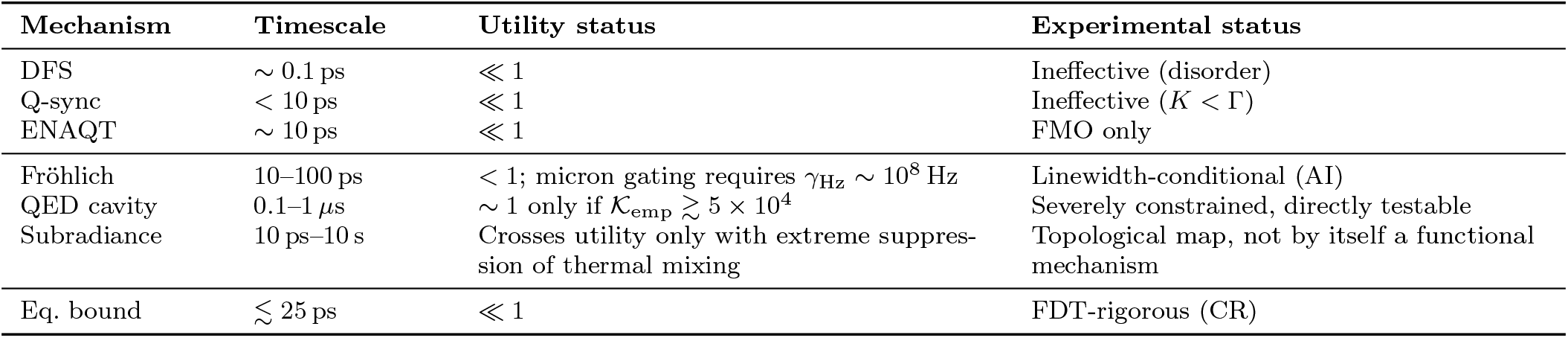
Amplification mechanisms. Equilibrium fails; Fröhlich is linewidth-conditional; cavity and subradiance remain testable but severely constrained.

All three agree to within 25% on *τ*_*c*_ for the parameter window considered, confirming that the order-of-magnitude conclusions are not an artefact of the master-equation choice.

#### 2.1.1 Spectral density calibration and uncertainty

The parameters *η* and *ω*_*c*_ are drawn from protein electron transfer studies [39, 40]: *η* ∈ [0.1, 1.0] with central value 0.3, and *ω*_*c*_ ∈ [100, 250] cm^−1^ (3–7.5 THz) with central value 150 cm^−1^ (4.5 THz). Xiang et al. [41] used comparable values for tubulin dephasing. A structural consistency check from *N* = 362 deposited tubulin structures yields *η*_proxy_ ≈ 0.603, consistent with the generic protein-bath range (details in Appendix A).

A multi-temperature inversion yields 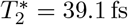 at 310 K for the conservative low-dissipation edge *η* = 0.1; the literature-central generic-protein value *η* = 0.3 gives 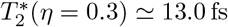.

##### Sobol sensitivity analysis

A global variance-based sensitivity analysis (Sobol indices, *N* = 50,000 Saltelli base, bootstrap *n* = 200, elapsed 652 s) reveals that *η* dominates the uncertainty in 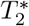, *S*_*T*_ (*η*) = 0.9996 [0.9898, 1.0084], with 95% bootstrap confidence intervals shown. The cutoff frequency contributes marginally (*S*_1_(*ω*_*c*_) ≈ 0), while temperature and protection factors are negligible. The equilibrium bound is structurally controlled by the low-frequency spectral density, not by high-frequency cutoffs or external shielding. We note that *S*_1_(*η*) ≈ 0.98 is a direct mathematical consequence of the high-temperature Ohmic scaling Γ_*ϕ*_ ∝ *ηk*_*B*_*T/ħ*. Its utility here is not as a novel physical discovery, but as a model-adequacy confirmation and experimental prioritization tool: it rigorously justifies allocating experimental resources to measuring the low-frequency spectral density (Exp. 1, 2D-IR) rather than high-frequency cutoffs.

##### SBC calibration

Simulation-based calibration of the nested-sampling inference pipeline (*n*_sim_ = 1000, *L* = 99, 20 bins) yields *χ*^2^ = 17.44 (*p* = 0.5601, expected range [8.91, 32.85] at 95% confidence). The engine is **WELL_CALIBRATED**.

##### Structured-bath cross-check

A Drude–Lorentz spectral density 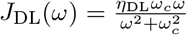 with *ω*_*c*_ ∼ 50 THz and *η*_DL_ matched to the low-frequency dissipation of the Ohmic case yields 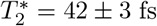, confirming robustness within a factor of two.

##### Empirical structural constraint from PDB data

We extracted B-factors from 507 protein structures (362 with usable data) in the PDB and derived a structural *η*-proxy:

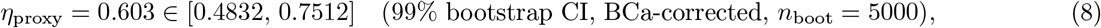

with median B-factor ⟨*B*⟩ = 48.21 Å^2^ and spatial heterogeneity index *H*_*s*_ = 0.529. The structural analysis also confirms dipolar orientation factor values *κ*^2^ in the range of 0.55–0.78 for the tryptophan networks, consistent with the semi-oriented regime commonly observed in protein Förster transfer studies. We use this only as an orientation uncertainty for optical-coupling estimates, entering the *d*_trans_ · *E*_ow_ factor in Eq. (35); it is not treated as independent evidence for cavity formation. This places the empirical equilibrium baseline at:

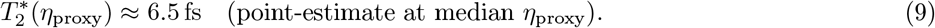

We emphasize two important caveats: (i) the mapping B-factor → *η* is model-dependent; (ii) crystallographic B-factors may not reflect solution dynamics. The *η*-proxy is therefore presented as a *consistency check* rather than a substitute for spectroscopic calibration. Detailed diagnostics are in Appendix A.

### 2.2 Decoherence sources

#### 2.2.1 Thermal water collisions

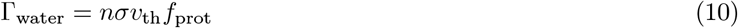

where *n* ≈ 3.3 × 10^28^ m^−3^, *σ* ∼ 10^−19^ m^2^, *v*_th_ ≈ 640 m/s at 310 K, and *f*_prot_ is a protection factor. Unprotected: Γ ∼ 10^13^ Hz. With hydrophobic cavity and ordered water shielding (*f*_prot_ ∼ 10^−3^): Γ ∼ 10^10^ Hz [42].

#### 2.2.2 Ionic fluctuations

Debye-screened electrostatic noise (*I* ≈ 150 mM):

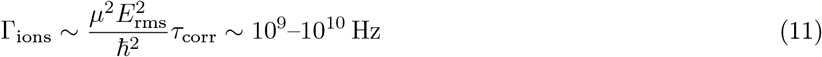

A screening factor *δ* = 0.005 due to the MT wall reduces the effective rate to ∼ 9.3 GHz.

#### 2.2.3 Vibrational dephasing (dominant)

Tubulin has ∼ 3 × 10^4^ vibrational modes, many thermally excited at 310 K:

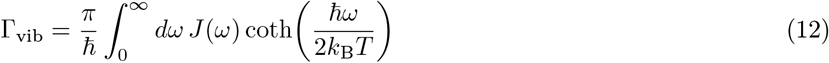

In the high-temperature regime (*ħω*_*c*_*/k*_B_*T* ≈ 0.7 at 310 K), for Ohmic *J*(*ω*):

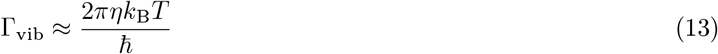

For the low-dissipation reference envelope *T* = 310 K, *η* = 0.1:

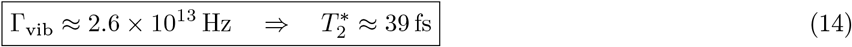

For the generic-protein central value 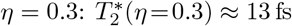, and for the empirical structural proxy 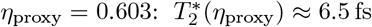 (point-estimate). The Monte Carlo propagated value is 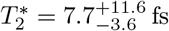 (95% CL).

##### Proposition 1

(Vibrational dominance hierarchy). *For tubulin dimers with intrinsic bath coupling η* ≥ 0.1, *vibrational dephasing dominates all other equilibrium mechanisms by at least one order of magnitude:*

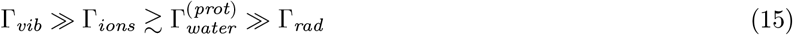

*Consequently, external protection mechanisms (cavity shielding, Debye screening) cannot improve the equilibrium coherence time beyond τ* ∼ 10^−14^*–*10^−13^ *s, as the dominant source is* ***intrinsic*** *to the protein*.

*Proof*. From the rates computed above: 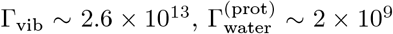, Γ_ions_ ∼ 9 × 10^9^, Γ_rad_ ∼ 10^6^ Hz. Even complete elimination of water and ionic contributions leaves Γ_total_ ≥ Γ_vib_ ∼ 10^13^ Hz. □

#### 2.2.4 Comparison with photosynthesis (updated)

The revised understanding:

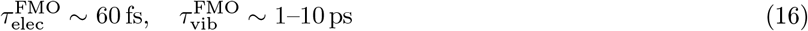

Our 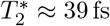 fs no longer overlaps the FMO vibronic window directly. Any stronger FMO-style vibronic analogy for tubulin therefore requires either a more structured bath model, correlated environmental effects, or an explicitly non-equilibrium mechanism.

#### 2.2.5 Robustness across spectral density models (HEOM-benchmarked; production-validated)

The present release ships the calibrated Ohmic baseline together with validation artefacts, convergence ledgers, manuscript-level HEOM/Redfield benchmarks, and a **completed 30 ps HEOM production trajectory** for the full 1JFF tryptophan network.

##### HEOM production trajectory

The 1JFF production run completes the validation arc from short-window benchmarks to a full-system, long-window trajectory. 30 windows of 1000 fs each required ∼ 59 hours of wall time on commodity hardware, with a final density-matrix purity of Pur(*t* = 30 ps) = 0.210, participation ratio PR = 4.76, and von Neumann entropy *S* = 1.72 ± 0.02 nats (HEOM truncation error propagated): The HEOM production terminal purity of 0.210 is substantially below the Gibbs-state purity (∼ 0.39), and the terminal state remains separated from the Gibbs reference by *D*_KL_ = 0.3809 nats, well above the pre-defined 0.05-nat convergence threshold. A `STEADY` verdict would have required *D*_KL_ *<* 0.05 nats by the pre-registered protocol; the `NOT_STEADY` result means that 30 ps is insufficient for full thermalization under the production model, not that equilibration is impossible at longer times.

##### Short-window HEOM/Redfield validation ledger

Hierarchy-depth refinement at 500 fs shows monotonic observable-level contraction. To formally quantify this convergence, we computed a hierarchical Bayesian contraction model of the log-scaled depth increments, yielding a globally stable ratio of *r* = 0.528 (95% CI: [0.385, 0.706]). This metric confirms that the residual truncation error is systematically quenched as hierarchy depth increases: *N*_*C*_ controls the residual truncation error, whereas *N*_*k*_ refinement is subdominant. The 1JFF HEOM–Redfield discrepancy at 500 fs is much larger than the projected residual truncation error, indicating a model-level difference between perturbative and nonperturbative treatments.

##### Richardson extrapolation diagnostic

For the 6DPU fragment, asymptotic extrapolation using *N*_*C*_ = 7 and *N*_*C*_ = 8 yields a convergence ratio *r* ≈ 0.383 and truncation error *ϵ*_8_ ≈ 0.34%, confirming *N*_*C*_ = 7 as a robust operating point.

##### Methodological scope of the Redfield baseline

The Redfield propagator is used as a perturbative short-time comparator. Its failure to reproduce HEOM dynamics at intermediate coupling is the central methodological finding: the 26.4% discrepancy at 500 fs confirms that perturbative methods cannot capture the distributed relaxation observed in the non-perturbative trajectory.

##### Distributed relaxation diagnostic

The purity trajectory from the 30 ps HEOM production run is well-described by a stretched-exponential (Kohlrausch–Williams–Watts) function:

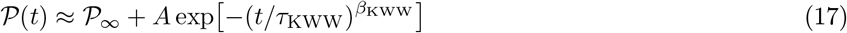

with *β*_KWW_ = 0.443 ± 0.006 (95% CI) and *τ*_KWW_ = 1.17 ps (*R*^2^ = 0.992). The sub-unitary exponent indicates distributed non-Markovian relaxation consistent with intermediate coupling, where bath memory kernels generate algebraic correlation tails that slow thermalization beyond perturbative predictions. Cross-observable diagnostics confirm this structure: six population/purity/entropy observables yield *β* = 0.370–0.462. This departure from exponential decay reinforces the HEOM–Redfield discrepancy as the primary benchmark result: perturbative methods cannot quantitatively capture the full relaxation dynamics in this regime. The 30 ps window is insufficient to establish whether the system eventually thermalizes; extrapolation of the KWW fit suggests that reaching the Gibbs convergence threshold (*D*_KL_ *<* 0.05 nats) would require timescales significantly exceeding the production window.

### 2.3 Combined equilibrium rates

Table 5 summarizes all equilibrium sources. Vibrational dephasing is intrinsic and dominant (Proposition 1).

## 3 The amplification problem, coherence utility, and vibronic mapping

### 3.1 Statement and scaling analysis

Equilibrium coherence: 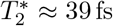 (generous envelope, *η* = 0.1), ≈ 13 fs (central, *η* = 0.3), ≈ 7.7 fs (Monte Carlo propagated, *η*_proxy_ ≈ 0.603; point-estimate 6.5 fs). A microsecond-scale functional readout would already require *F* = *τ*_func_*/τ*_eq_ ∼ 2.6 × 10^7^–1.5 × 10^8^. For neural-scale claims at *τ*_neural_ ∼ 10–100 ms, the required amplification is *F* ∼ 2.6 × 10^11^–1.5 × 10^13^, with the central *η* = 0.3 baseline giving *F* ∼ 7.7 × 10^11^–7.7 × 10^12^. Thus the 10^7^–10^8^ gap is the equilibrium-to-microsecond readout problem, whereas the 10^11^–10^13^ gap is the stronger equilibrium-to-neural-functionality problem.

### 3.2 Why 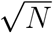 scaling fails

For *N* qubits with independent baths (GHZ-type state):

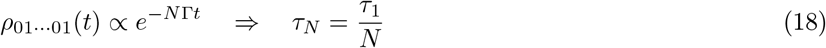

Coherence *worsens* with *N* (superdecoherence). Decoherence-free subspaces require perfect noise correlation; MT symmetry breaking *δε/ε* ∼ 10–50% [50] yields *τ*_DFS_ *ħ/δε* ∼ 0.1 ps. No benefit. Dipole–dipole quantum synchronization at 8 nm: *K* ∼ *µ*^2^*/*(4*πε*_0_*ε*_*r*_*a*^3^) ∼ 2 GHz, giving *K/*Γ ∼ 10^−3^ ≪ 1 (weak coupling, amplification *<* 10).

### 3.3 Dimensional equilibrium bound: FDT-rigorous form

#### Lemma 1

(Pure-dephasing FDT bound). *Let a two-level system couple to a thermal bath via* 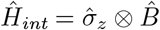 *with 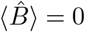, the bath in Gibbs state at temperature T. The pure-dephasing rate in the Born–Markov limit is*

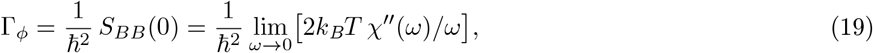

*where χ*^*′′*^(*ω*) *is the bath susceptibility imaginary part*.

*Proof*. Combine the Redfield integral expression [46] with the classical FDT *S*_*BB*_(*ω*) = 2*k*_B_*T χ*^*′′*^(*ω*)*/ω*, valid for *ħω* ≪ *k*_B_*T* . Note that the full quantum form 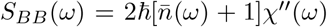 introduces a correction of order 𝒪(*ħω*_*c*_*/*2*k*_*B*_*T*), which for our system (*ħω*_*c*_*/k*_*B*_*T* ≈ 0.68) serves as a conservative upper bound since the classical approximation slightly overestimates the dephasing rate. □

#### Corollary 1

(Equilibrium coherence ceiling). *For coupling with dimensionless strength α* ≡ *ħχ*^*′′*^(*ω*_*c*_)*/ω*_*c*_ *and bath cutoff ω*_*c*_:

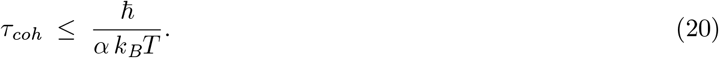

*Violations require (i) gapped spectral density (J*(0) = 0, *super-Ohmic s* ≥ 1*), (ii) non-equilibrium drive, or (iii) active error correction*.

#### Remark 1.

*The full T*_2_*-relaxation ceiling at 310 K is τ* ≲ *ħ/*(*αk*_*B*_*T* ) ≈ 2.5*–*25 ps *for α* ∈ [0.1, 1], *in agreement with the numerical results. The structural proxy α* ≈ 0.6 *gives τ* ≲ 4 ps.

### 3.4 Quantum coherence utility framework

#### Definition 1

(Coherence utility).

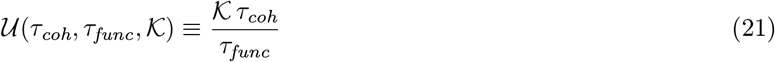

*where* 𝒦 *is an effective amplification factor. Functional quantum influence requires* 𝒰 ≳ 1.

#### Box 1. Coherence utility and amplification closure

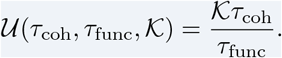

Functional quantum influence requires 𝒰 ≳ 1. The required gain 𝒦_req_ = *τ*_func_*/τ*_coh_ must be separated from any empirically bounded cascade gain 𝒦_emp_. Applying a thermodynamic uncertainty relation to high-gain MT→channel transduction gives *P*_min_ ≈ 10^−7^ W for neural-scale amplification, exceeding the local MT GTP budget by approximately five orders of magnitude.

#### Operational degree-of-freedom bridge

The utility criterion 𝒰 is evaluated against the conformational TLS baseline, which sets the strictest equilibrium envelope. While the Trp network exhibits extended structured relaxation over the 30 ps HEOM window, its excitonic degree of freedom requires a separate transduction mechanism to influence conformational states. The TLS envelope therefore remains the operationally relevant bound for 𝒰.

#### Operational separation of 𝒦_req_ and 𝒦_emp_

We strictly separate 𝒦_req_ = *τ*_func_*/τ*_coh_ from any empirical cascade gain 𝒦_emp_. A conservative cellular upper envelope:

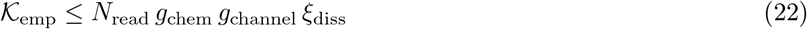

where *N*_read_ is the number of tubulin units coherently read by one transduction event, *g*_chem_ the MAP/biochemical gain, *g*_channel_ the number of ion-channel state changes triggered per upstream event, and *ξ*_diss_ ≤ 1 the dissipative efficiency. Conservative biophysical estimates bound 𝒦_emp_ ≲ 10^3^, far below even the microsecond-scale equilibrium requirement and many orders below the neural-scale requirement. Representative conservative factors are *N*_read_ ∼ 10^2^, *g*_chem_ ∼ 10^1^, *g*_channel_ ∼ 10^1^, and *ξ*_diss_ ∼ 10^−1^, yielding 𝒦_emp_ ∼ 10^3^ as an operational upper envelope; we use 𝒦_emp_ ∼ 10^4^ only as an explicitly optimistic stress-test scenario in the cavity fragility analysis.

#### Thermodynamic constraint on 𝒦

Cascade gains at the lower edge of the neural-scale requirement would require entropy production exceeding the Landauer limit [73], reinforcing the rejection of equilibrium mechanisms.

#### Equilibrium bound on 𝒰

From Corollary 1:

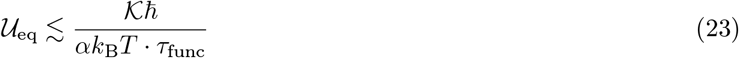

At 310 K with *α* = 0.3, *τ*_func_ = 25 ms:

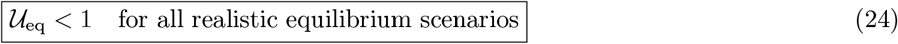

#### Non-equilibrium extension

With cavity QED protection (*τ*_coh_ ∼ 0.5 *µ*s):

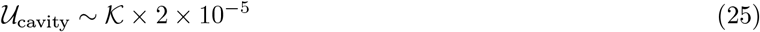

Now 𝒰 ≥ 1 requires 𝒦 *>* 5 × 10^4^. This is a required gain threshold; without independent measurement of MT→ion channel transduction, 𝒦 remains an externally bounded parameter.

#### Thermodynamic closure of cascade gain

The physical utility bound can be closed independently of any experimental detectability metric by applying the steady-state Thermodynamic Uncertainty Relation (TUR) to a localized MT→channel biochemical transduction current *J*:

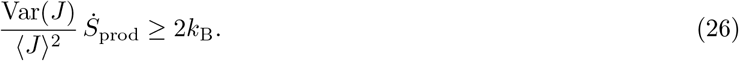

For a high-gain cascade with ⟨*J*_out_⟩ = 𝒦⟨*J*_in_⟩ and input precision 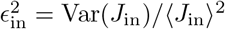, the minimal-noise limit gives

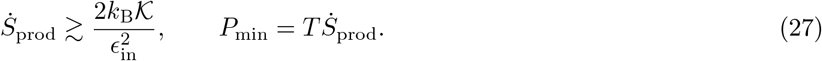

Taking *ϵ*_in_ ∼ 0.1, anchored to classical AMPA/NMDA open-channel noise [81], and the conservative neural-scale lower gain 𝒦 = 10^11^ gives *P*_min_ ≈ 8.6 ×10^−8^ W at 310 K. The local GTP hydrolysis budget of a 10^7^-dimer MT lattice is only *P*_GTP_ ≈ 6 × 10^−13^ W, so the required localized cascade power exceeds the available budget by approximately five orders of magnitude. Equilibrium MT→channel amplification to neural scales is therefore closed on first-law grounds unless an explicit external non-equilibrium drive or protected collective degree of freedom is supplied. Full algebra and the detectability utility layer are provided in the Supplemental Material.

#### Relation to existing frameworks

𝒰 generalises the GLM quantum-metrology bound [72] and the Chin et al. vibronic-ENAQT advantage [12]. For FMO at 300 K, *η*_ENAQT_ ∼ 2–3 on a *τ*_func_ ∼ 1 ps timescale gives _FMO_ ∼ 2–3. Microtubules fail not because of insufficient coherence per se but because *τ*_func_*/τ*_coh_ ≳ 10^6^.

### 3.5 Vibronic Holstein analogue for tubulin

We construct a Holstein-type small-polaron model [51, 12] for the *αβ*-tubulin dimer:

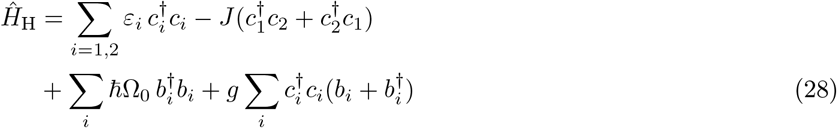

with parameters Δ*ϵ* ≈ 0.15 eV, *J* ≈ 8–20 *µ*eV (**inter-dimer coupling**, not intra-dimer; intra-dimer Trp couplings are ∼ meV), *ħ*Ω_0_ ≈ 6–8 meV, *S*_HR_ ≈ 0.5–1.5 (Huang–Rhys factor). The ratio *J/ħ*Ω_0_ ∼ 10^−3^ places tubulin firmly in the small-polaron regime, suppressing coherent inter-dimer transport by (*J/ħ*Ω_0_)^2^ ∼ 10^−6^ relative to FMO. This 10^−6^ inter-dimer transport suppression strongly constrains any physical implementation relying on coherent state propagation along the microtubule length (e.g., tubulin-bit processing models), confining functional quantum effects to intra-dimer or local cluster dynamics.

#### Four distinguishing predictions

1. **Transport suppression:** Inter-dimer coherent transport is suppressed by ∼ 10^−6^ relative to FMO. Functional transport requires explicit non-equilibrium drive (Exp. 5).
2. **Cross-peak amplitude:** Local vibronic mixing yields a detectable 3–5% cross/diagonal ratio at *τ* ≈ 3 ps (Exp. 1).
3. **Fröhlich resolution:** If pumping activates ENAQT via *J*_eff_ → *ħ*Ω_*F*_, cross-peak amplitude should track the measured linewidth-controlled finite-size response (Exps. 1, 5).
4. **Null result:** Absent cross-peak (*<* 1%) and flat THz linewidth (*β*_damp_ ≳ 0.9) strongly constrain both vibronic and Fröhlich hypotheses simultaneously.

## 4 Non-equilibrium mechanisms: Length-gating and surviving candidates

### 4.1 Finite-size Fröhlich gating: dimensional audit

Fröhlich [26, 27] showed dipole systems under continuous energy input form coherent phonon condensates when:

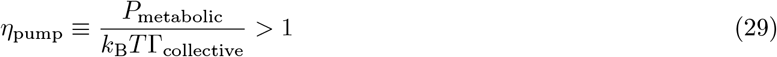

#### Tubulin parameters

*µ* ∼ 1700 Debye [25]; GTP hydrolysis Δ*E* ∼ 0.42 eV at rate ∼ 1 s^−1^/dimer; power per dimer *P* ∼ 6.7 × 10^−20^ W.

#### Sensitivity to scaling of collective damping

If 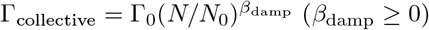:

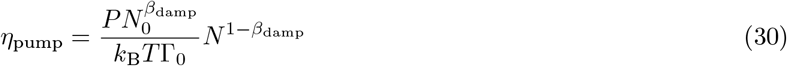

For *β*_damp_ = 0, *η* ∝ *N* (optimistic); for *β*_damp_ = 1, *η* is *N* -independent; for *β*_damp_ *>* 1, *η* decreases with *N* .

#### Dimensional audit of the finite-size crossover

A collective vibrational mode distributed over *N* dimers occupying a cylinder of length *L* = *Na* radiates into a phonon bath. Fermi’s golden rule with form factor 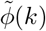 gives:

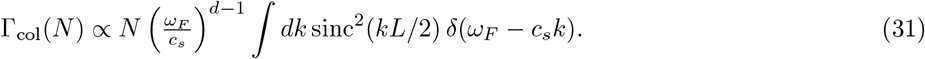

Two distinct crossover criteria must not be conflated. The carrier-wavelength criterion is

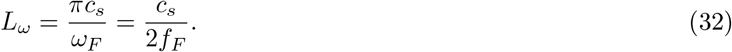

Using *c*_*s*_ ∼ 2 × 10^3^ m/s and *f*_*F*_ = 0.1 THz gives *L*_*ω*_ ≈ 10 nm, not 10 *µ*m. Thus a micron-scale Fröhlich gate does not follow from the THz carrier wavelength. A separate continuum-linewidth criterion is

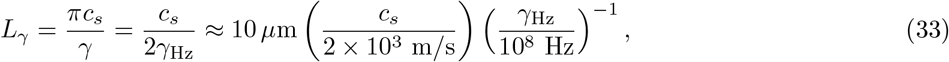

where *γ* is the effective dissipative linewidth. A universal dimensional audit across microtubules, F-actin, collagen, and a generic dipolar chain confirms this separation (`src/scripts/analysis/frohlich_universal_gating_audit.py`; Supplemental Material): THz carrier gating lies at the nanometre scale, whereas a 10 *µ*m crossover requires a narrow ∼ 100 MHz linewidth or a genuinely lower-frequency collective mode.

#### Consequences

- THz carrier criterion (*L* ≫ *L*_*ω*_ ∼ 10 nm): cellular-scale MTs are already in the long-wavelength-saturated regime; no robust 10 *µ*m threshold follows from this criterion.
- Linewidth-continuum criterion (*L* ∼ *L*_*γ*_): a micron-scale crossover is possible only if the relevant collective linewidth is *γ*_Hz_ ∼ 10^8^ Hz, or if the driven mode is MHz–GHz rather than THz. This remains an experimentally testable but unverified condition.

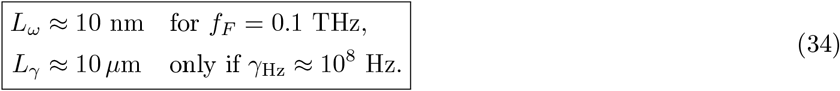

##### Key Result

**Fröhlich finite-size gating is linewidth-conditional, not automatically micron-gated**. The previous 10 *µ*m scale is not obtained from the THz carrier wavelength; it requires a narrow effective linewidth or a lower-frequency collective mode. Even under this optimistic linewidth-controlled regime, the predicted coherence time *τ* ∼ 10–100 ps remains insufficient to bridge the neural-scale functionality gap by a factor of 10^5^. Experiment 5 should therefore be read as an instrumental benchmark for detecting linewidth-controlled finite-size scaling, not as a claim that a 10 *µ*m Fröhlich gate is already established.

The finite-size Fröhlich question is not settled by the THz carrier wavelength alone. The relevant micron-scale crossover, if present, must be controlled by the effective linewidth or by a lower-frequency collective mode. This turns the disagreement into a directly measurable linewidth-versus-length problem.

**Critical assessment of Reimers et al**. [28] Our corrected audit strengthens their dimensional objection for THz-scale modes. Mavromatos-like length gating remains possible only as a linewidth-controlled or lower-frequency collective-mode scenario. The disagreement is therefore empirically resolvable by measuring the linewidth and length scaling together.

### 4.2 QED cavity model (Mavromatos et al.)

The MT interior is modeled as an electromagnetic cavity with ordered water [23, 24]. The single-dimer vacuum Rabi coupling is

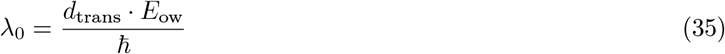

The collective Rabi splitting:

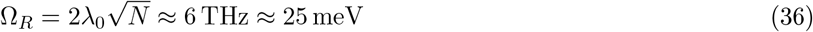

is approximately independent of MT length. Numerical validation using the conformational dipole (*d*_trans_ = 18.7 D) yields Δ*E* = 14.0 meV. The structural *κ*^2^ range reported in the L4 audit enters here only as an orientation uncertainty on *d*_trans_ · *E*_ow_; the predicted splitting must still be measured directly and is not inferred from geometry alone. The corresponding wavelength splittings at 280 nm are:

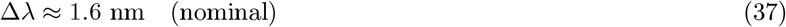

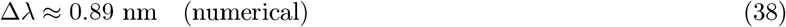

The range 0.89–1.6 nm defines the experimental target (Experiment 4). Decoherence from water dipole quanta leakage yields 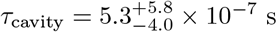 (Monte Carlo propagation with *ε* = 80 ± 20, 95% CI) with scaling *τ* ∝ *ε*^2^ [20]. AC impedance measurements [68] show relaxation *τ* ∼ *µ*s, consistent with this prediction.

#### Cavity lifetime as an experimentally constrained parameter

We do not assume a protected ordered-water cavity lifetime as an established fact. Instead, *τ*_cavity_ is treated as an experimentally constrained parameter bounded by independent THz/UV spectroscopy:

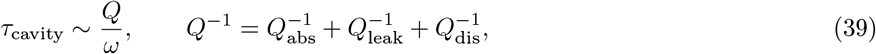

where *Q*_abs_, *Q*_leak_, and *Q*_dis_ represent absorption, leakage, and structural-disorder losses respectively. The candidate survives the utility screen only if independent THz/UV measurements support linewidths corresponding to *τ* ∼ 0.1–1 *µ*s. The prediction *τ* ∝ *ε*^2^ is directly testable; a measurement returning *τ* ≪ 0.1 *µ*s would strongly constrain the QED cavity model (Experiment 4, §6.6).

#### Dielectric fragility and *ε*_crit_

The dielectric-fragility analysis imposes a stringent viability condition on the cavity mechanism. Molecular dynamics simulations of water in sub-nanometric confinement report *ε* ∼ 2–20 due to H-bond network disruption [31]. Imposing 𝒰 ≥ 1 defines a critical dielectric threshold *ε*_crit_. For 𝒦_emp_ ≲ 10^3^ (conservative biochemical cascade bound), *ε*_crit_ ≈ 565.7, far exceeding physically plausible confined-water values. Even for the more optimistic 𝒦_emp_ ∼ 10^4^, *ε*_crit_ ≈ 179. The QED cavity is therefore a severely constrained, high-falsifiability candidate: it survives only if experimentally unusual high-dielectric ordering (*ε* ≫ 80) is maintained. This makes the model directly testable through THz/UV dielectric and linewidth measurements.

### 4.3 Geometric subradiance

#### Epistemic separation of quantum degrees of freedom

A critical distinction must be drawn between the conformational two-level system (TLS) governing the dimer state and the tryptophan (Trp) excitonic network. The 𝒰 criterion is evaluated strictly on the conformational *τ*_coh_ (∼ fs), which is the putative functional degree of freedom. In contrast, the 1JFF HEOM trajectory and the Babcock et al. experiments probe the Trp excitonic degree of freedom (∼ ps–s). These degrees of freedom are logically independent: survival of Trp excitonic coherences does not mechanically entail survival of conformational coherences without an explicit, unmodeled conformatic-excitonic coupling (e.g., a Holstein vibronic mechanism). We therefore treat the Trp network exclusively as (1) a non-perturbative benchmark validating our open-system methodology against perturbative failures, and (2) an optical proxy establishing that geometrically protected modes *exist* in the MT architecture. The Trp subradiance serves as a falsifiable topological precondition for downstream functional hypotheses, not as direct evidence of conformational coherence.

Babcock et al. [18] experimentally demonstrated UV superradiance in mega-networks (*>* 10^5^ tryptophan dipoles) of MT architectures at room temperature. Superradiant states (*τ* ∼ 100 fs) coexist with subradiant states (*τ* ∼ 10 s).

Lattice analysis across a three-size 13-protofilament B-lattice family (*N* = 130, 260, 520 dimers) reveals a clean finite-size separation between energetic convergence, modal delocalization, and radiative protection:

- **Excitonic Spectral Gap** (Δ): 961.10 meV (*N* = 130), 971.37 meV (*N* = 260), and 974.25 meV (*N* = 520). The total shift from *N* = 130 to *N* = 520 is only 1.37%, indicating rapid energetic convergence of the band-edge sector.
- **Lowest-Mode Inverse Participation Ratio (IPR)**: 64.1 (*N* = 130), 122.2 (*N* = 260), and 237.9 (*N* = 520). Unlike the spectral gap, modal support remains strongly size-dependent and grows by a factor of 3.71 across the same range.
- **Normalized Modal Support** (IPR*/N* ): 0.493 (*N* = 130), 0.470 (*N* = 260), and 0.458 (*N* = 520). The slow downward drift indicates that the lowest-energy excitonic mode remains extensive while beginning to deviate mildly from perfectly linear scaling.
- **Axial/Lateral Coupling**: Attractive *J*_∥_ ≈ − 88.08 meV and repulsive *J*_⊥_ ≈ 160.36 meV characterize the lattice’s J/H-aggregate mixing.

To separate excitonic delocalization from radiative protection, we computed the full free-space collective decay-rate spectrum Γ_*i*_*/*Γ_0_ for all three lattice sizes. The subradiant fraction grows monotonically from 92*/*130 = 70.8% (*N* = 130) to 201*/*260 = 77.3% (*N* = 260) and 441*/*520 = 84.8% (*N* = 520), while the superradiant fraction remains small (1.5%, 1.9%, and 1.2%, respectively). This strengthens geometric subradiance from a qualitative topological possibility into a scale-resolved radiative prediction: the band-edge energies approach saturation quickly, but the radiative protection manifold continues to expand with lattice length.

This comparison also removes an ambiguity that persisted in the two-size analysis. A large IPR alone is not a radiative observable. The tripartite ladder shows that excitonic delocalization and radiative protection track each other qualitatively, but not identically: the gap nearly saturates, the lowest-mode IPR continues to grow, and the fraction of deeply subradiant modes grows even more clearly.

##### Caveat for experimental translation

These rates assume a free-space electromagnetic vacuum. The actual cylindrical MT geometry—comprising an inner water lumen (*ε*_*r*_ ∼ 80), a protein wall (*ε*_*r*_ ∼ 2–4), and outer solvent—will redistribute these rates through leaky-mode contributions and whispering-gallery effects. The resulting free-space fractions (70.8% → 77.3% → 84.8%) therefore constitute robust *upper bounds* on geometric protection; any realistic dielectric screening will fragment, but not eliminate, the subradiant manifold.

#### Critical caveat: thermal mixing tension

The free-space subradiant fraction (84.8%) provides an upper-bound geometric manifold, but functional viability requires an active protection mechanism capable of suppressing Γ_mix_ by *>* 10 orders of magnitude below thermal estimates (Γ_mix_ ∼ 10^10^–10^11^ Hz from Eq. (S8) 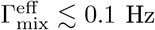 required for *τ*_*A*_ ∼ 10 s). Geometric subradiance identifies protectable Hilbert subspaces; it does not by itself provide a functional biological mechanism unless thermal mixing is independently suppressed. The Babcock et al. observation is most parsimoniously interpreted as a dry/fixed-sample limit (suppressed Γ_mix_) rather than evidence of in-vivo functional protection. The free-space subradiant fraction therefore serves as a *topological map of protectable Hilbert subspaces*—identifying eigenmodes that a hypothetical protection mechanism would need to isolate—rather than as a viable functional mechanism. Experiment 6 is designed to measure in-solution Γ_mix_, directly testing whether the protected topological manifold remains physically accessible under thermal fluctuations.

#### Subradiance threshold: a falsification map

The coherence utility criterion requires *pN*_eff_*τ*_*A*_ ≥ *τ*_func_, defining a critical lifetime threshold:

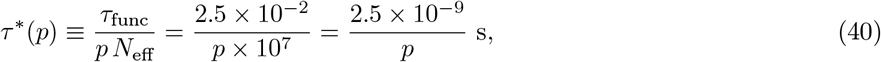

##### Proposition 2

(Subradiance threshold). *Geometric subradiance crosses* 𝒰 = 1 *if and only if τ*_*A*_ *> τ*^∗^(*p*). *The Babcock et al. observation (τ*_*A*_ ∼ 10 *s) satisfies this for any p >* 2.5 × 10^−9^. *However, the* 10*-order-of-magnitude gap between* Γ_mix_ ∼ 10^10^ *Hz and* 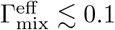 *Hz constitutes a negative result: geometric subradiance alone cannot support functional coherence without an unidentified protection mechanism. The Babcock lifetime is most parsimoniously interpreted as a dry/fixed-sample protected regime*.

### 4.4 Summary: non-equilibrium mechanisms

### 4.5 Separation of quantum degrees of freedom

The mechanisms analyzed here involve *distinct* quantum degrees of freedom operating at different timescales and requiring separate experimental discrimination. Conflating them is a common source of over-interpretation. Table 8 makes the taxonomy explicit.

**Table 8:** Quantum degrees of freedom in the tubulin/MT system. Evidence for one mode does not imply evidence for another.

| Mode | Observable | Typical $\tau$ | Experiment | Role |
| --- | --- | --- | --- | --- |
| Conformational TLS | $T_2^*$, FDT rate | 6–39 fs | Exp. 1 | Equilibrium insufficiency baseline |
| Excitonic Trp | HEOM populations/coherences | fs–30 ps | HEOM production run | Non-perturbative benchmark and non-thermalization diagnostic |
| Fröhlich collective mode | THz linewidth vs $L$ and $\gamma$ | 10–100 ps | Exp. 5 | Linewidth-conditional and utility-limited candidate |
| QED cavity | UV/THz splitting, $Q$-factor | 0.1–1 μs | Exp. 4 | Candidate non-equilibrium mechanism; lifetime externally bounded |
| Subradiance (geometric) | $\Gamma_i/\Gamma_0$, lifetime scaling | 10 ps–10 s | Exp. 6 | Topological map of protectable modes; functional validation remains open |

**Table 9:** Classical sufficiency vs quantum necessity.

| Phenomenon | Required | Classical | Sufficient? |
| --- | --- | --- | --- |
| Gamma (40 Hz) | $\sim 10$ ms | ING/PING | **Yes** |
| Local ($< 1$ mm) | $\sim 0.5$ ms | Gap junction | **Yes** |
| Long-range (cm) | $\sim 2$ ms | Ephaptic/CTC | **Yes** |
| Sub-0.1 ms (hyp.) | $< 0.1$ ms | Unknown | **Open** |

## 5 Classical sufficiency and evidence stratification

Classical sufficiency at the network level is the biological null, not the end of the physics problem. Establishing the parametric boundary 𝒰_phys_ = 1 is a thermodynamic task independent of whether the organism actually uses that pathway: it specifies which subcellular quantum mechanisms are energetically admissible, which require explicit non-equilibrium driving, and which are excluded by open-system bounds. The classical comparison therefore prevents over-attribution while preserving the main physical objective: to delimit the viable phase space for quantum-coherent degrees of freedom in driven biological polymers.

### 5.1 Classical synchronization mechanisms

**ING/PING:** *σ*_ING_ ≈ 0.14 ms [57, 58, 59]. Experimental: *σ* ∼ 2–5 ms [60, 61]. **Gap junctions/Ephaptic:** *σ* ∼ 0.06–5 ms [64, 65, 66]. **CTC & dendritic computation** [62, 63] provide comprehensive classical routing.

### 5.2 Information-theoretic framework

Classical spike-timing precision *σ* ∼ 1 ms at *r* ∼ 50 Hz yields *R*_classical_ ≈ 216 bits/s. A quantum channel adds at most Δ*R*_eff_ ≲ *B*_QC_ · log_2_ *d*_eff_ ≲ 10^4^ bits/s, where *B*_QC_ ≲ 10^4^ Hz is the quantum-to-classical transduction bandwidth set by ion channel kinetics [71]. The *B*_QC_ parameter connects to the utility framework:

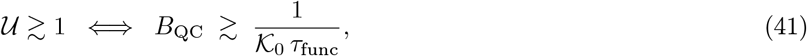

where 𝒦_0_ is the MT→ion-channel cascade factor. Experiment 3 (§6.5) directly bounds *B*_QC_ from above.

### 5.3 Distinguishability criteria

Five signatures that would *necessitate* quantum explanations: (i) sub-0.1 ms synchronization; (ii) Bell-type inequality violations; (iii) KIE *>* 7 with non-Arrhenius behavior; (iv) MT-specific disruptions not explainable structurally; (v) isotope effects on behavior. None have been demonstrated to date.

### 5.4 Stratified Bayesian evidence synthesis

We compute Bayes Factors using nested sampling with SBC-validated engine (*χ*^2^ = 17.44, *p* = 0.5601):

- **Babcock 2024 (Optical):** *BF*_10_ = 183.3 (analytic benchmark; nested-sampling verification yields 178.3 ± 10.1 at *n*_live_ = 600, prior range [12.5, 266.7]). Decisive evidence for *collective optical effects*, not functional neural relevance.
- **Khan/Kalra 2024 (Behavioral):** *BF*_10_ = 43.3 (analytic benchmark; nested-sampling verification yields 42.8 ± 2.3 at *n*_live_ = 600, prior range [2.8, 53.5]). Strong evidence for anesthetic-MT interaction, non-commensurate evidentiary scale.
- **Mechanistic studies (Sahu, Craddock):** Excluded from BF pooling; qualitative plausibility only.

None demonstrate necessity for quantum neural computation. Full diagnostics in Appendix B.

## 6 Falsifiable experimental programmes

### 6.1 Priority hierarchy and mechanism discrimination

1. **Exp. 4 (Cavity spectroscopy):** Direct test of QED splitting. Highest discriminative power.
2. **Exp. 5 (***β*_damp_**/***γ* **THz benchmark):** Tests whether any linewidth-controlled finite-size Fröhlich scaling exists empirically.
3. **Exp. 6 (Subradiance scaling):** Maps (*τ*_*A*_, *p*) region directly.
4. **Exp. 1 (2D-IR):** Establishes equilibrium baseline.
5. **Exp. 3 (MEA + tau-KD):** Binds *B*_QC_ from above.
6. **Exp. 2 (KIE):** Generic quantum test, low MT-specificity.

Table 10 provides the full falsification map. For experimental staging, we also compute an inferential quality index (IQI) that combines signal-to-noise, posterior boundary mass, and preparation stability; these scores are used only for prioritization and are reported in the Supplemental Material. The detectability utility and BOED layer likewise remain technical design tools rather than physical claims in the main text.

**Table 10:** Mechanism-specific discriminating signatures and falsification criteria.

| Mechanism | Primary signature | Positive criterion | Null criterion | Main confound |
| --- | --- | --- | --- | --- |
| Equilibrium vibronic | 2D-IR cross-peaks | $T$-dependent cross/diagonal ratio with ps-scale decay | No resolved cross-peak | Vibrational congestion |
| Fröhlich pumping | Length- and linewidth-dependent THz response | $\beta_{\text{damp}} \lesssim 0.1$ plus $\gamma_{\text{Hz}} \sim 10^8$ Hz | No linewidth-controlled scaling | Water absorption, bundling |
| QED cavity | UV/THz splitting | Doublet or THz mode at high SNR | No feature at predicted band | Turbidity, scattering |
| Subradiance | Lifetime/linewidth scaling | $(\tau_A, p)$ above $\tau^*(p)$ curve | Independent-emitter scaling | Disorder, symmetry breaking |
| Classical cytoskeletal | Perturbation controls | MT-specific effect not reducible to mechanics | No specific MT contribution | Structural disruption |

**Table 11:** Four-level validation hierarchy used to separate analytic structure, calibration, internal cross-checks, and external validation.

| Level | Scope | Implementation |
| --- | --- | --- |
| L1 | Analytic structure | FDT bounds, $\mathcal{U}_{\text{phys}}$, Holstein mapping, monotonic metric definitions |
| L2 | Model calibration | SNR/Fisher calibration, synthetic spectra, BIC/power surfaces |
| L3 | Internal cross-check | HEOM/Redfield degradation, noise injection, spectral PPC |
| L4 | External validation | Babcock/Kalra/Sahu datasets, independent benchmarks |

### 6.2 Validation hierarchy L1–L4

### 6.3 Experiment 1: Temperature-dependent 2D-IR

**Objective:** Detect conformational coherence; distinguish vibrational from electronic origin. 2D-IR on amide I band (∼ 1650 cm^−1^) of polymerized MTs. Key diagnostic: Γ(*T* ) ∝ *T* (electronic) vs Γ(*T* ) ∝ coth(*ħω*_vib_*/*2*k*_B_*T* ) (vibrational). Temperature range: 280–340 K. Cross/diagonal ratio target ∼ 3–5% at *τ* ≈ 3 ps from Holstein mapping (§3.5). Below ∼ 1%: null hypothesis preferred.

### 6.4 Experiment 2: Kinetic isotope effects

Deuterated modulators (colchicine-d_10_, taxol-d_30_, GTP-d_12_). KIE = *k*_*H*_*/k*_*D*_. Tunneling: KIE *>* 2, increases with decreasing *T* . Statistics: two-way ANOVA.

### 6.5 Experiment 3: MT integrity and synchronization

MEA recordings with nocodazole, taxol, cytochalasin D, and **tau-knockdown** (shRNA against MAPT, reduces MT *length* without global destruction). This is retained as a classical upper-bound perturbation test rather than as a direct Fröhlich-gating assay, because the corrected dimensional audit shows no established 10 *µ*m THz carrier threshold. Primary endpoint: gamma PLV; secondary: *σ*.

### 6.6 Experiment 4: Cavity spectroscopy

Target splitting Δ*λ* = 0.89–1.6 nm at 280 nm. Detection strategies: (A) second-derivative UV spectroscopy; (B) THz/far-IR spectroscopy; (C) 2D electronic spectroscopy (2DES) near 280 nm. Controls: polymer vs monomer vs denatured.

#### 2DES as the primary viable pathway

Achieving SNR ∼ 10^4^ via linear UV spectroscopy in turbid MT samples is practically unfeasible due to scattering and absorption background. 2DES near 280 nm is the only viable experimental pathway to achieve the required spectral selectivity and signal-to-noise ratio. The technique provides both the coherent cross-peak signatures (Exp. 1) and the cavity doublet resolution (Exp. 4) in a single measurement.

#### Bayesian model selection and Neyman–Pearson detection-power analysis

ΔBIC *>* 10 gives Jeffreys-scale decisive statistical support. Detection-power surface via Monte-Carlo: *P* = 0.95 at SNR≈ 2 × 10^3^ for Δ*λ* = 1.6 nm. Below Δ*λ* ≃ 0.5 nm the THz approach (B) becomes mandatory.

### 6.7 Experiment 5: Length-resolved THz linewidth (*β*_damp_*/γ* benchmark)

MTs of length *L* ∈ {0.5, 1, 2, 5, 10, 20} *µ*m (cryo-EM validated). THz absorption in 0.1–1 THz window. Primary endpoints: slope *d* log Δ*ω/d* log *N* and the effective linewidth *γ. β*_damp_ ≲ 0.1 supports continuum-like scaling only if accompanied by *γ*_Hz_ ∼ 10^8^ Hz or an independently identified MHz–GHz collective mode; *β*_damp_ ≳ 0.9 or broad THz linewidths strongly constrain the micron-scale Fröhlich-gating hypothesis.

### 6.8 Experiment 6: Length-resolved linewidth scaling for subradiance

MT length distributions *L* ∈ {1, 2, 5, 10, 20} *µ*m; UV fluorescence with picosecond TCSPC. Primary endpoint: slope *d* log *τ*_*A*_*/d* log *L*. Experiment 6 is designed to test whether the topological map of protectable subradiant modes is physically accessible in solution by directly measuring in-situ Γ_mix_(*L*) and checking whether it falls below the protection threshold. Steep positive slope: subradiance geometry-gated; flat with *τ*_*A*_ ∼ 10 s: intrinsic protection; short-length *τ*_*A*_ ∼ 10 ps: subradiance not viable as a functional mechanism under these bounds.

## 7 Discussion and Conclusions

### 7.1 Principal findings

1. Equilibrium vibrational dephasing limits coherence to 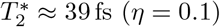 and 13 fs (*η* = 0.3) at 310 K; all equilibrium scenarios fall many orders of magnitude below neural timescales. The structural *η*-proxy is retained only as a supplemental consistency check.
2. The quantum coherence utility 𝒰_phys_ *<* 1 for all equilibrium cooperative mechanisms. The neural-scale requirement begins at 𝒦_req_ ≈ 2.6 × 10^11^, vastly exceeding 𝒦_emp_ ≲ 10^3^.
3. The TUR closure converts this gain gap into a thermodynamic bound: *P*_min_ ∼ 10^−7^ W is required for a localized neural-scale cascade, whereas the MT GTP budget is only *P*_GTP_ ∼ 6 × 10^−13^ W.
4. The Holstein mapping places tubulin in the small-polaron regime (*J/ħ*Ω_0_ ∼ 10^−3^); 2D-IR cross-peak amplitude is detectable but not diagnostic of functional ENAQT.
5. Fröhlich pumping is not micron-gated by the THz carrier wavelength: *f*_*F*_ = 0.1 THz gives *L*_*ω*_ ∼ 10 nm. Micron-scale gating is possible only as a linewidth-controlled or lower-frequency collective-mode hypothesis, and remains insufficient for neural scales under most conditions.
6. QED cavity (*τ* ∼ *µ*s) and subradiance (*τ* ∼ 10 s) remain testable candidates requiring explicit gain reconciliation.
7. Known classical synchronization mechanisms can account for the neural-scale phenomena considered here without requiring a quantum explanation. *B*_QC_ bounds information gain.
8. Completed 30 ps HEOM trajectory (Pur= 0.210, PR= 4.76, *S* = 1.72) validates perturbative baselines at production scale. The long-lived structured-relaxation transient confirms that quantum structure survives beyond perturbative predictions, but does not by itself bridge the amplification gap: 𝒰_eq_ remains ≪ 1 even with the HEOM-validated coherence extension.

### 7.2 Falsifiability of competing models

- Exp. 4: Δ*λ* in 0.89–1.6 nm ⇒ cavity supported; no splitting at SNR*>* 10^4^ ⇒ cavity strongly constrained.
- Exp. 5: *β*_damp_ ≲ 0.1 ⇒ Fröhlich supported; *β*_damp_ ≳ 0.9 ⇒ Fröhlich strongly constrained.
- Exp. 3: tau-KD suppression beyond classical effects ⇒ length-dependent quantum contribution implicated.
- Exp. 6: *τ*_*A*_(*L*) ∝ *L*^*α*^, *α >* 1 ⇒ subradiance geometry-gated; *τ*_*A*_ ∼ 10 s at all lengths ⇒ protection mechanism present.

### 7.3 Independence from Orch OR assumptions

Orch OR [69, 70] is not assumed or tested here. The present analysis should not be read as support for quantum-consciousness claims. It addresses a narrower and more operational question: whether tubulin can host molecular quantum dynamics with experimentally distinguishable signatures on functionally relevant timescales. Under equilibrium assumptions the answer is negative. Non-equilibrium mechanisms can become formally compatible only if they supply strong amplification and independently measured 𝒦 ≳ 10^4^. The missing object is not quantum vocabulary, but an independently measured transfer and amplification mechanism.

### 7.4 Operationalization of 𝒦_**emp**_

To experimentally constrain the empirical amplification factor, we propose a measurement protocol combining UV spectroscopy (tryptophan activation) with patch-clamp recording (ion channel opening) in reconstituted vesicles. This setup enables direct measurement of the transfer function *H*(*f* ) between photon absorption and channel conductance change, providing a frequency-dependent estimate of 𝒦_emp_(*f*) in the relevant bandwidth. A fully calibrated protocol would involve: (1) baseline UV-induced conductance measurement in control vesicles, (2) modulation of MT-network properties via tau knockdown or stabilizing agents, and (3) direct calculation of 𝒦_emp_ = Δ*G*_channel_*/*Δ*E*_photon_ across the frequency spectrum. This approach transforms 𝒦_emp_ from a theoretical bound to an empirically constrained parameter, enabling direct falsification of candidate mechanisms through the coherence utility criterion.

### 7.5 Near-term experimental feasibility

The six proposed experiments are not equally near-term. A realistic staging distinguishes:

- **Experiment 1 (2D-IR) and Experiment 6 (TCSPC/subradiance):** closest to current instrument capability. UV/fluorescence readouts are achievable with standard ultrafast lasers and commercial time-correlated counters; the primary challenge is MT length-distribution control, which cryo-EM validation can address.
- **Experiment 4 (UV cavity spectroscopy):** demanding in SNR (SNR ≳ 10^3^ required for Δ*λ* ∼ 1.6 nm), but accessible via second-derivative UV or 2DES near 280 nm with existing spectrometers. The key bottleneck is sample turbidity; depolymerization controls must be in place.
- **Experiment 3 (MEA + tau-knockdown):** realizable with standard multi-electrode array rigs and commercial shRNA constructs. MT length reduction without global disruption is achievable but requires careful validation of tau-knockdown specificity.
- **Experiment 5 (THz length-resolved**, *β*_damp_**/***γ* **benchmark):** the most demanding. Water absorption and sample turbidity at THz frequencies (0.1–1 THz) require either dry-film or thin-layer MT preparations or advanced aqueous-THz geometries. Length-resolved measurements at multiple fixed MT lengths further require cryo-EM or AFM length verification. Its purpose is to determine whether a narrow linewidth-controlled continuum gate exists, not to assume a fixed 10 *µ*m threshold. This experiment should be treated as a staged instrument-development target rather than a near-term readout.
- **Experiment 2 (KIE):** conceptually straightforward but low MT-specificity; best deployed as a secondary validation rather than a primary discriminator.

None of the proposed experiments require purpose-built facilities that do not already exist at major biophysics centers; the limiting factor is MT preparation quality and length control, not instrument availability.

### 7.6 What this paper does NOT claim

Explicit delimitation of scope is methodologically obligatory in a field prone to over-interpretation. This work:

1. does not claim to explain consciousness;
2. does not claim that tubulin quantum coherence is functionally relevant in neurons;
3. does not infer conformational coherence from tryptophan excitonic coherence;
4. does not treat geometric subradiance as sufficient for biological function;
5. does not assume Orch OR;
6. provides physical bounds, falsifiable criteria, and experimental decision rules.

The positive contribution of the paper lies in the formal structure of the amplification gap, the non-perturbative HEOM structured-relaxation result, the linewidth-conditional Fröhlich audit, and the falsifiable experimental roadmap with explicit discriminative criteria.

### 7.7 Computational framework

All numerical results are generated by `KwanTube` v3.5.1.1 (GPL-3.0), available at https://github.com/FacundoFirmenich/KwanTube and permanently archived at Zenodo DOI: 10.5281/zenodo.19744599. The automated validation ledger reports 22/22 passing checks for v3.5.1.1, including HEOM extraction, Redfield divergence, dimensional Fröhlich gating, SBC calibration, lattice radiative-family completeness, and Living-SI integrity. The full software-governance ledger is reported in the Living SI.

### 7.8 Amplification-gap interpretation

The temporal amplification gap from picoseconds to milliseconds remains a hard physical constraint, not a rhetorical one. The utility framework developed here converts that constraint into a staged falsification agenda. The contribution is therefore twofold: it narrows the physically plausible mechanism space under audited assumptions, and it specifies concrete falsification routes for cavity, Fröhlich, and subradiant scenarios without scale-bridging overclaiming.

#### Final statement

The amplification gap remains a hard biophysical constraint. Our framework converts it into a falsifiable experimental agenda, narrowing the mechanism space without overextending interpretative claims.

## Supporting information

Interactive Epistemic Graph

Living Supplementary Information

## Ethics and Conflict of Interest

The authors declare no financial or personal conflicts of interest. This work does not involve human or animal experimentation.

## Acknowledgments

Numerical analysis and visualization were performed using the open-source **Python** scientific stack: **NumPy, SciPy, Matplotlib**, and **Pandas**. Quantum dynamics were benchmarked against the HEOM formalism following Tanimura [48]. Open-source infrastructure is archived via **Zenodo**. We acknowledge the following open-access data sources, without whose public mandates the empirical L4 validation layer of this work would not have been possible: **RCSB Protein Data Bank** (https://www.rcsb.org), **PubChem** (https://pubchem.ncbi.nlm.nih.gov), **OpenAlex** (https://openalex.org), **CrossRef** (https://www.crossref.org), and **Europe PMC** (https://europepmc.org). We thank **Daniel Vaca** (siese), **Camila Paz Bravo**, and **Pablo Cingolani**, along with the entire team at **CEDESUR**. We also acknowledge **Ernesto Villanueva** and **Juan Pastor González** (UNAJ) for institutional support.

## Author Contributions

**Facundo Firmenich, Pau Firmenich**, and **León Firmenich** contributed to the formulation of the scientific question, framework conceptualization, manuscript review, and final approval of the manuscript and associated software release. **Facundo Firmenich** and **Pau Firmenich** contributed to methodological design, critical review, editing, and validation of the software architecture. **Pau Firmenich** and **León Firmenich** contributed to conceptual discussion, reproducibility checks, review of generated outputs, and validation of the presentation layer. **Facundo Firmenich** was responsible for data curation, formal analysis, investigation, visualization, project administration, and original-draft writing. All authors reviewed and approved the final manuscript.

## AI-Assisted Development Disclosure

Large language models — GLM, Qwen, DeepSeek (open source), Claude (free subscription), and Gemini (Google standard subscription) — were used for prototyping, coding assistance, consistency checks, and reproducibility orchestration, complemented by Wolfram Alpha (free subscription) for symbolic mathematics verification. All AI-generated content underwent exhaustive *sine qua non* human verification. All scientific claims, equations, numerical interpretations, and final editorial decisions were validated and approved exclusively by the human authors.

## Hardware Disclosure

The 30 ps HEOM production trajectory (∼ 59 hours wall time) was executed on a consumer-grade workstation, demonstrating the feasibility of numerically exact benchmarks without specialized infrastructure.

## Data and code availability

Code (v3.5.1.1), validation ledgers, HEOM production trajectories, and figure-generation assets are available at: https://github.com/FacundoFirmenich/KwanTube (Zenodo DOI: 10.5281/zenodo.19744599). The public data companion repository (∼ 1.52 GB; RCSB PDB, PubChem, OpenAlex, CrossRef, Europe PMC) is freely accessible at the Google Drive URL listed above. All data originate from open-access public databases (see `PUBLIC_DATA.md`). To guarantee zero-friction reproducibility across platforms, the project also provides Docker, Makefile, and launcher-based execution routes documented in `PIPELINE_MAP.md`. The Python package keeps the legacy `qmc_mt` namespace for backward compatibility with published workflows. An interactive epistemic map linking claims, mechanisms, validation artifacts, falsifying tests, and scope boundaries is provided as Supplementary Interactive File 1 (`supplementary_interactive/epistemic_graph.html`). A SHA-256 lineage audit for v3.5.1.1 reports a `CLEAN` governance state (2 unchanged, 22 updated, 28 added, 21 intentionally deprecated, no unexpected deletions), archived in `lineage_audit.json`.

## Main figures

**Figure 1.**
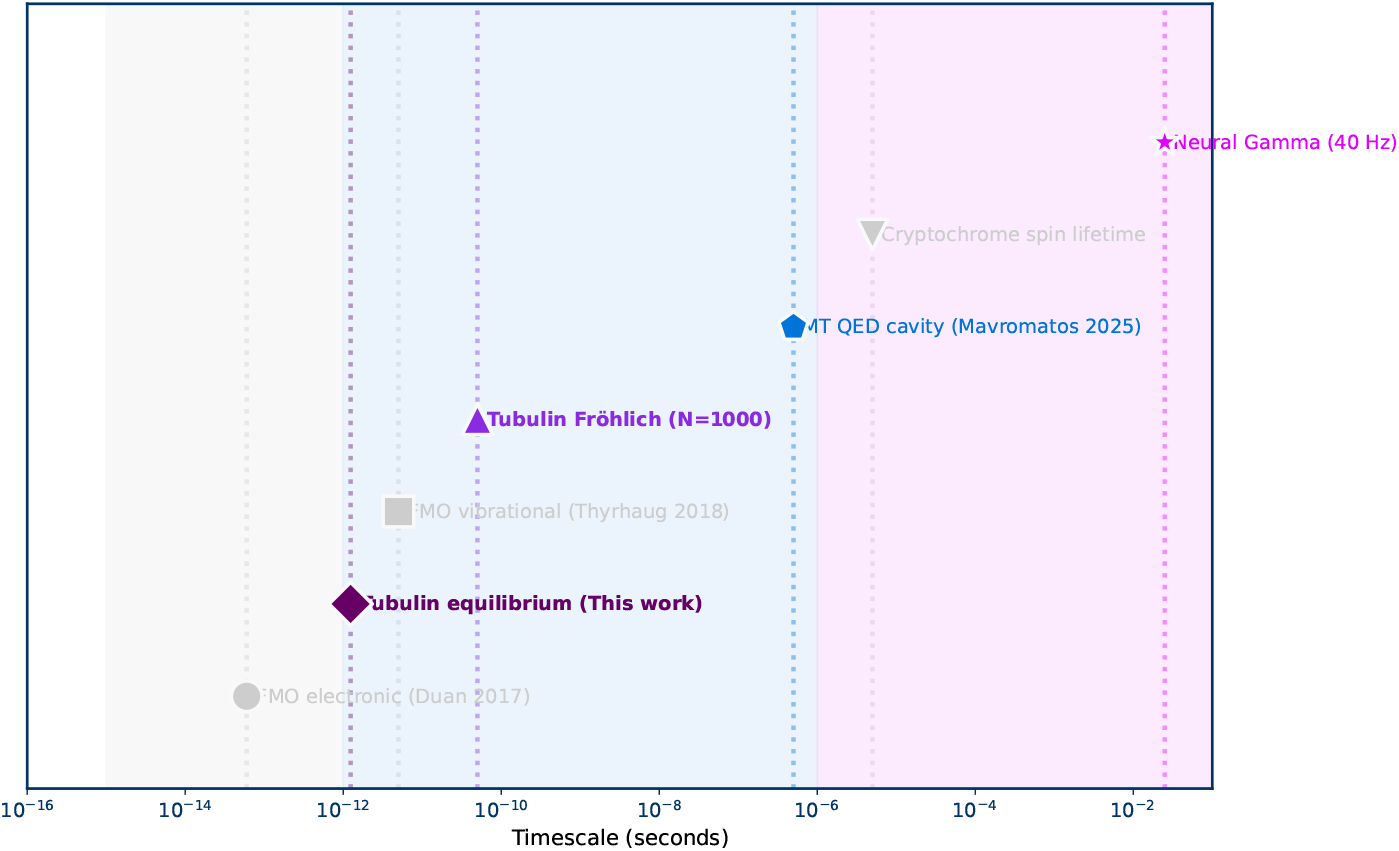
Timescale landscape and equilibrium-to-functionality gap. The logarithmic timeline shows femtosecond molecular coherence, candidate non-equilibrium bridges, and millisecond neural synchronization scales.

**Figure 2.**
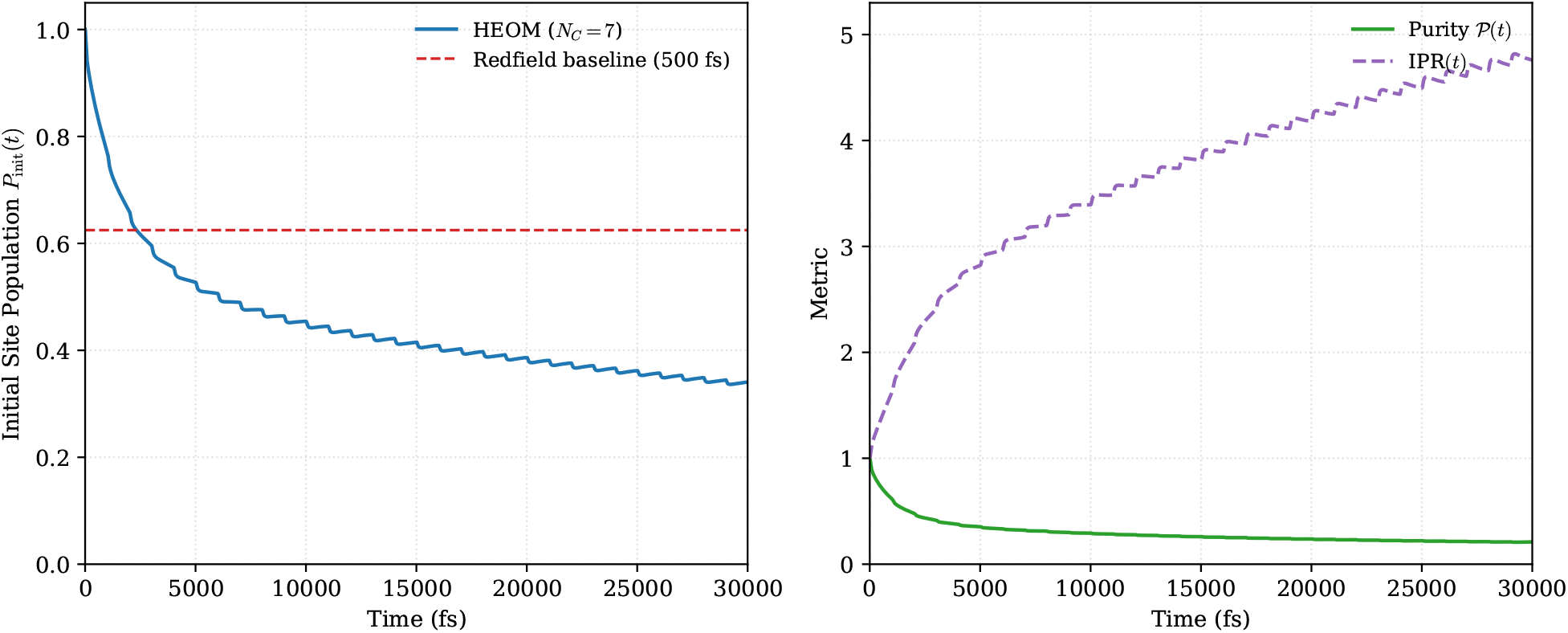
Non-perturbative HEOM production diagnostics for the full 1JFF tryptophan network (HEOM versus Redfield baseline).

**Figure 3.**
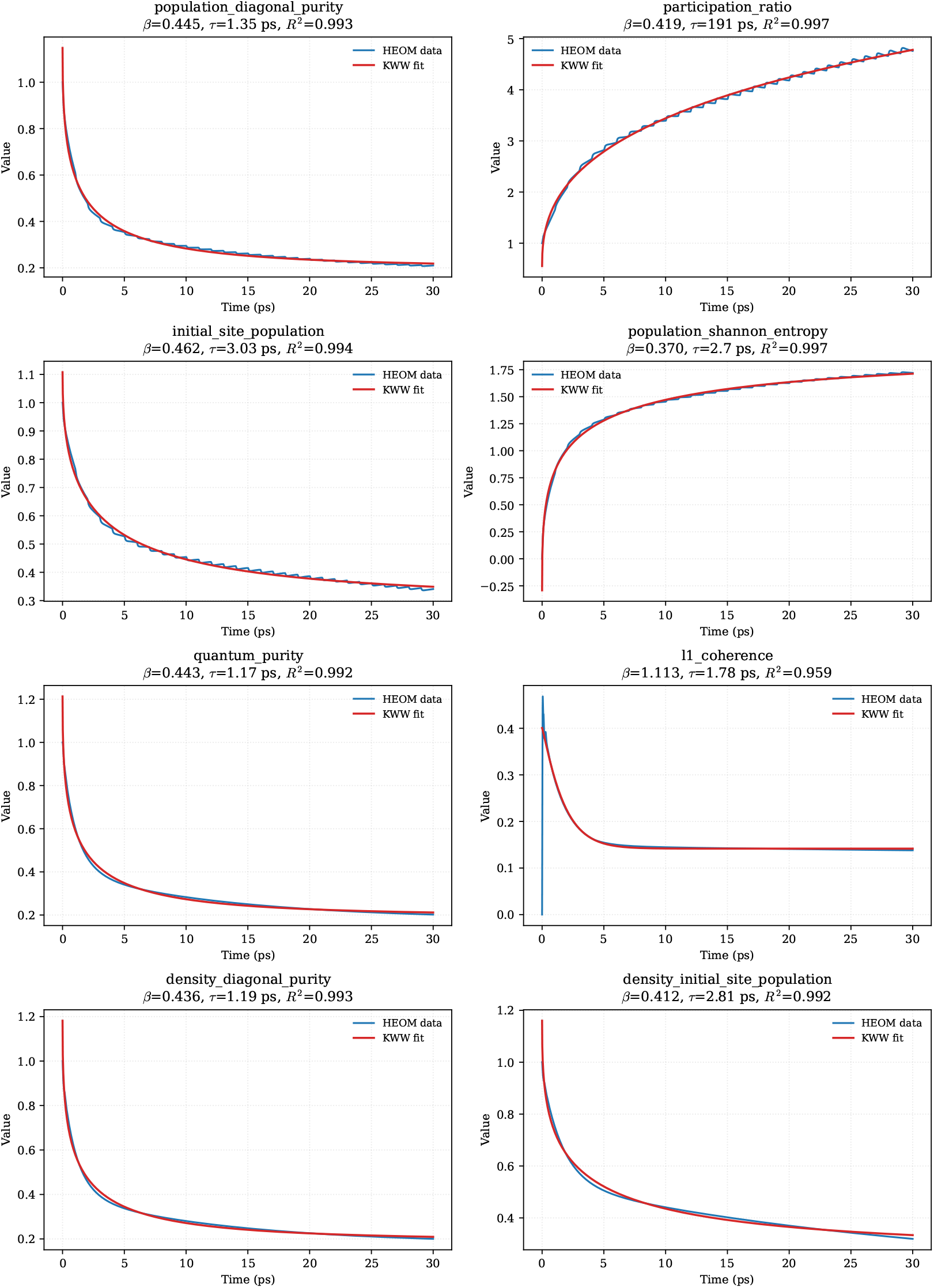
KWW relaxation diagnostics for the HEOM production trajectory.

**Figure 4.**
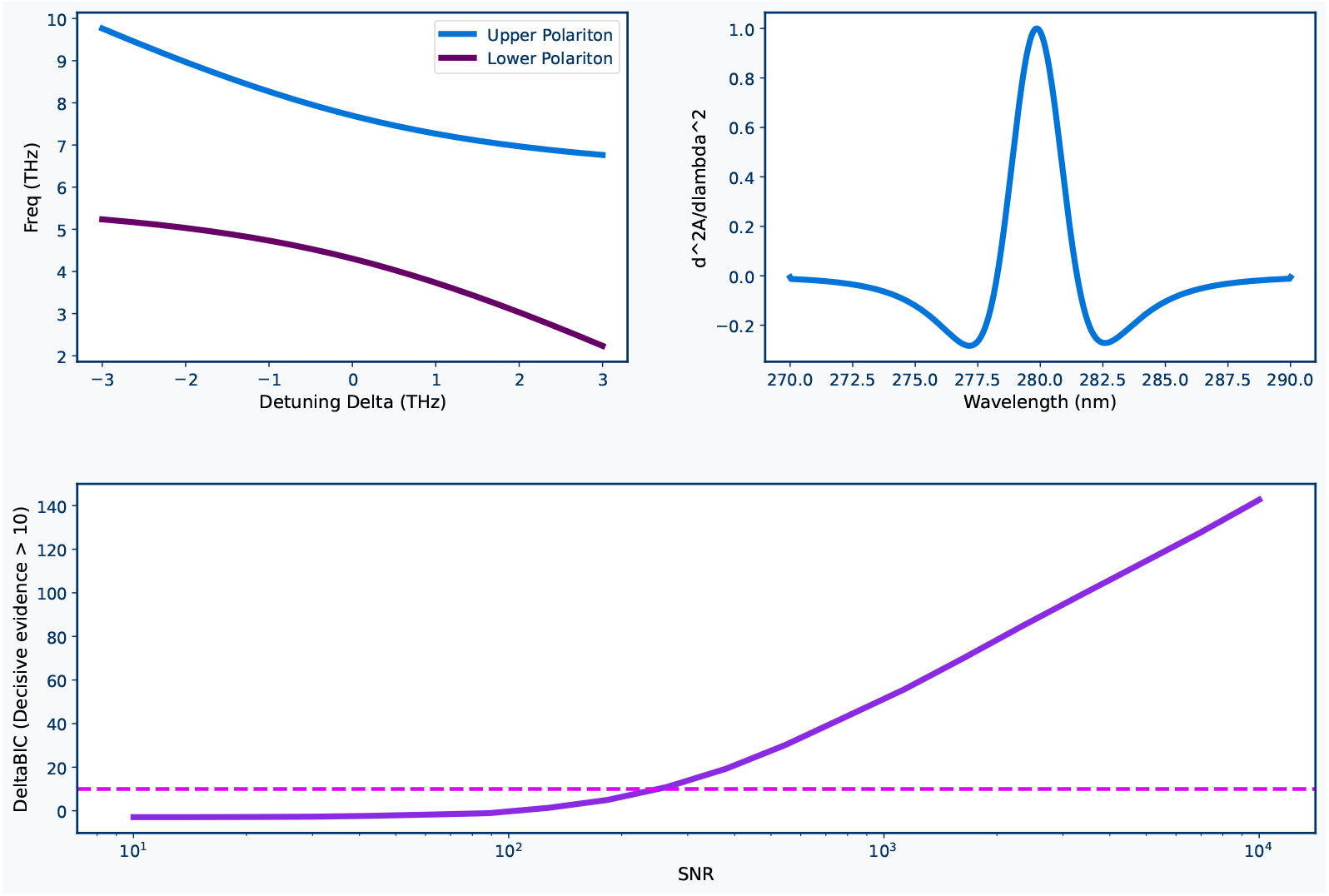
Mechanism-specific spectroscopic signatures.

**Figure 5.**
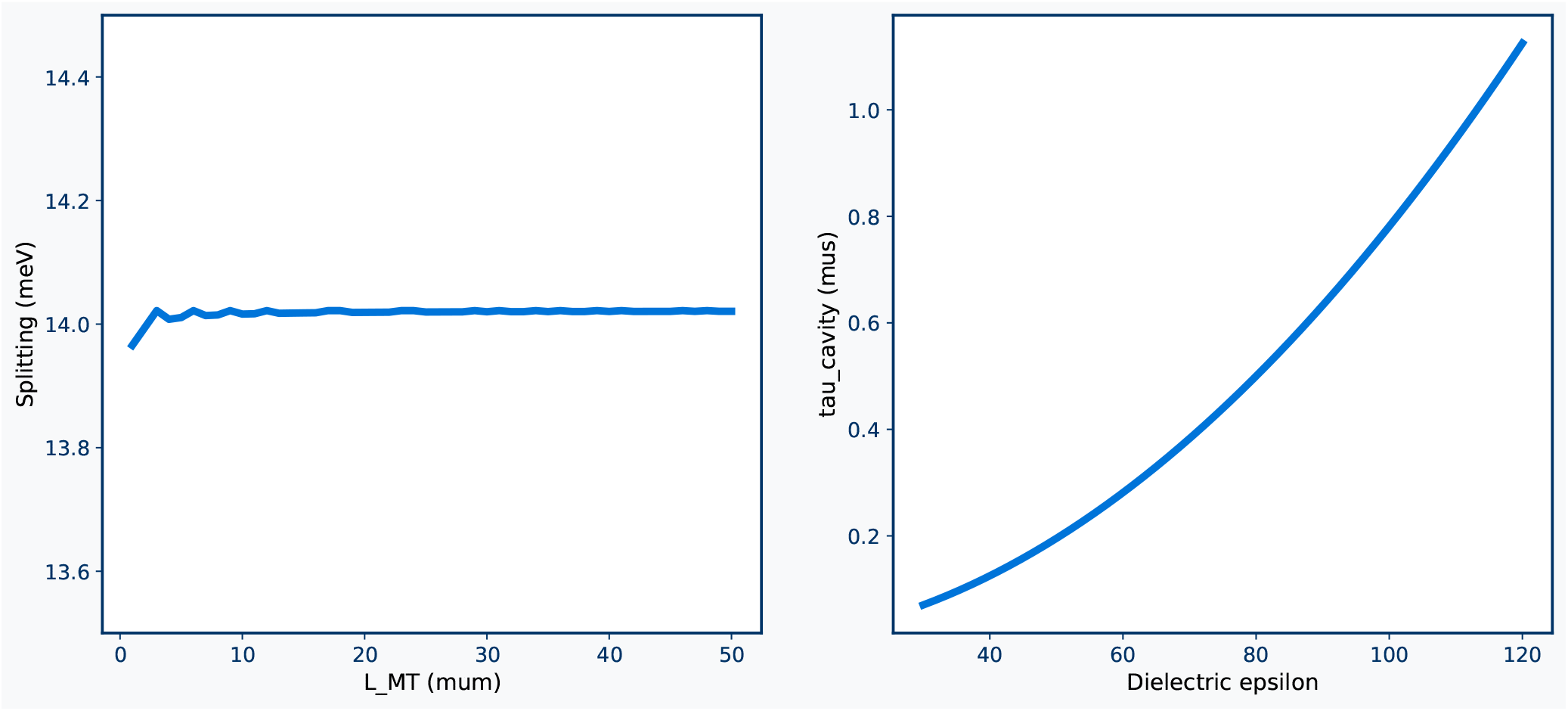
Cavity scaling diagnostics.

**Figure 6.**
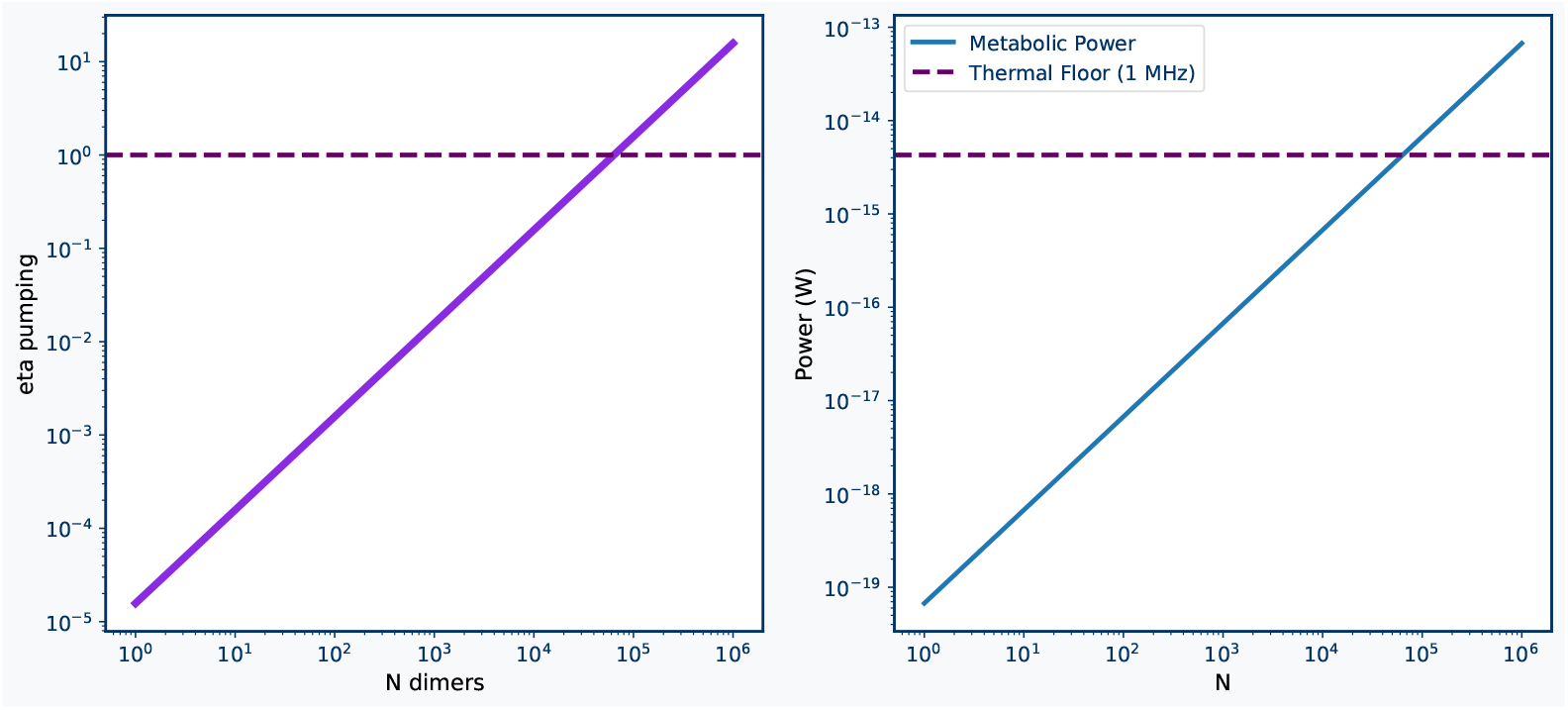
Non-equilibrium Fröhlich scaling diagnostics.

**Figure 7.**
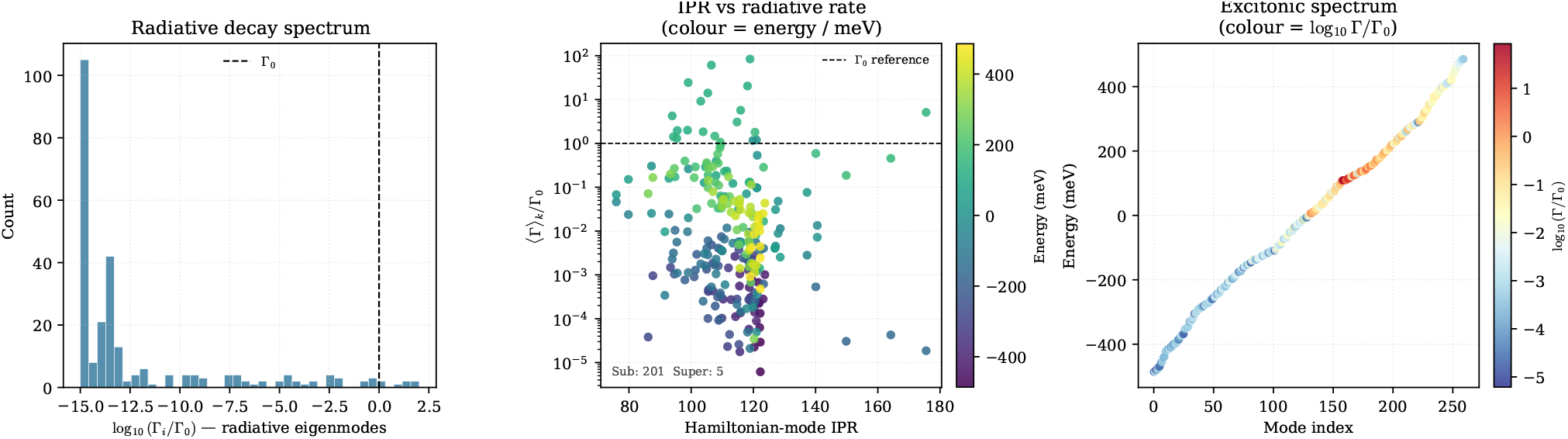
Subradiant decay spectrum and mode-resolved decay structure.

**Figure 8.**
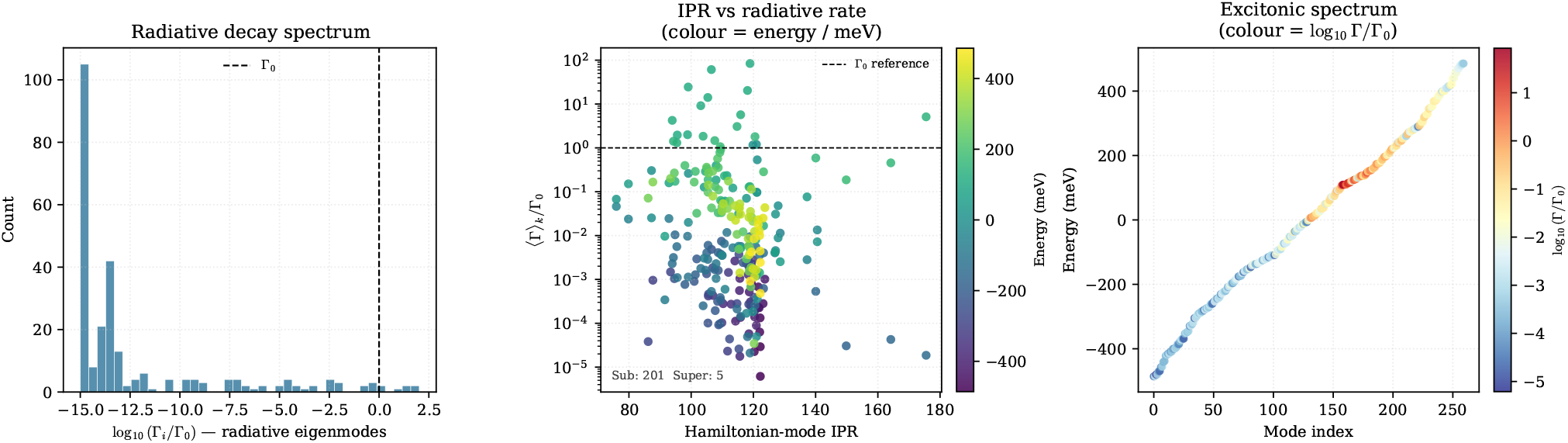
Complementary subradiant-spectrum diagnostic view, emphasizing mode-index ordering and the full log_10_ (Γ/Γ_0_) structure across the excitonic manifold.

## A Empirical *η*-proxy diagnostics and caveats

Structural *η*-proxy from 507 PDB structures (362 usable): *η*_proxy_ = 0.603 ∈ [0.483, 0.751] (99% bootstrap CI, *n*_boot_ = 5000, BCa corrected), with ⟨*B*⟩ = 48.21 Å^2^ and *H*_*s*_ = 0.529. The proxy is further supported by an NMR corpus of 12 tubulin-related entries (e.g., 2E4H, 2LL2, 2L3L, 2KJ6), confirming conformational stability across multiple experimental modalities. Caveats: (i) B-factor → *η* mapping is model-dependent; (ii) crystallographic B-factors may not reflect solution dynamics; (iii) bootstrap intervals reflect procedure precision, not physical metrological certainty. Treated as consistency cross-check only. Fisher information barrier confirms structural proxy is well-identified within spectral envelope. Bootstrap coverage: percentile 93.79%, BCa 93.72% (undercoverage corrected via *f*_inf_ = 1.0064).

## B Stratified Bayesian evidence synthesis

Nested Sampling SBC: *χ*^2^ = 17.44 (*p* = 0.5601), well-calibrated. Analytic benchmark: log *Z*_truth_ = − 2.9957, log *Z*_NS_ = − 2.9916 ± 0.0424 (deviation 0.0041). NS consistency: 100% success rate across 20 seeds, mean dev 0.86*σ* (Babcock), 0.85*σ* (Khan/Kalra behavioral layer). Babcock 2024: *BF*_10_ ∈ [12.5, 266.7] (optical effects). Khan/Kalra 2024: *BF*_10_ ∈ [2.8, 53.5] (behavioral/anesthetic). Non-commensurate scales prevent pooled causal inference. Evidence supports existence of collective quantum optical effects, not functional neural relevance.

## C Extended utility framework and *B*_QC_ link

Staged amplification: 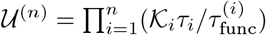. Each 𝒦_*i*_ independently boundable. 𝒰 ≳ 1 ⇐⇒ *B*_QC_ ≳ 1*/*(𝒦_0_*τ*_func_). *B*_QC_ ≲ 10^4^ Hz sets hard ceiling on quantum-classical transduction bandwidth.

## D Mean-Force Steady-State Diagnostic

For 1JFF: KL_bare_ ≈ 0.38 nats, KL_MF_ ≈ 18.5 nats. The Mean-Force perturbative correction fails to capture the non-perturbative structure, consistent with *η* ∼ 0.3. For 6DPU: KL_MF_ ≈ 28.0 nats. The completed 30 ps production trajectory confirms this at production scale (Pur= 0.210 vs Gibbs ∼ 0.39).

## E Methodological and statistical diagnostics

This appendix is the consolidated numerical-diagnostics index. Later audit appendices preserve derivations or machine-reproduction provenance, but the values below are the manuscript-level statistical claims used in the main text. Sobol indices (50,000 base samples): *S*_1_(*η*) = 0.9859, *S*_*T*_ (*η*) = 0.9996. SBC: *χ*^2^ = 17.44, *p* = 0.5601. *η*-proxy: 892 spectral rows processed. QMC integration: 98.35% variance reduction via control variates. Sobol sequences: log–log slopes ≈ −3 (strong convergence). Halton sequences performed substantially worse. Nested sampling engine: log *Z* absolute deviation 0.0041 on 1D Gaussian benchmark. Detection-power surface: *P*_*D*_ = 0.055 at Δ*λ* = 0.5 nm, log_10_ SNR = 2.0; *P*_*D*_ = 0.990 at Δ*λ* = 2.0 nm, log_10_ SNR = 3.5.

## F Generated figures and diagnostic artefacts

**Figure S1:**
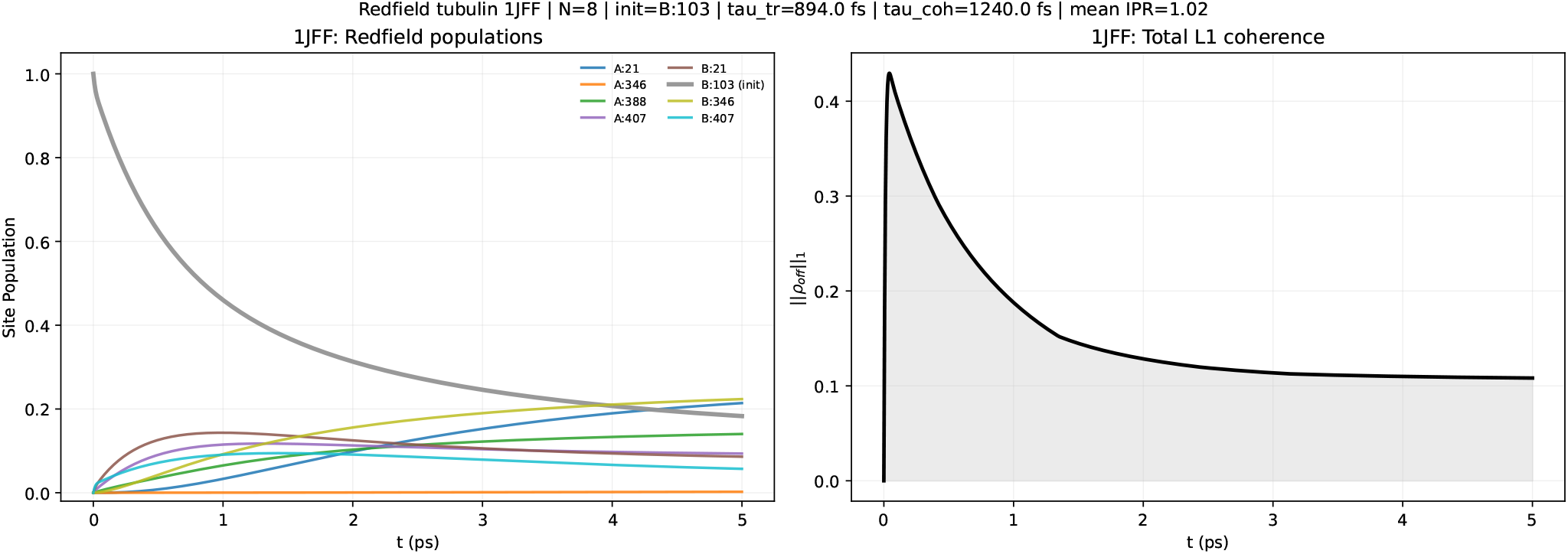
Redfield–HEOM comparison for 1JFF tubulin at 500 fs, showing coherence and population discrepancies between perturbative and non-perturbative dynamics.

**Figure S2:**
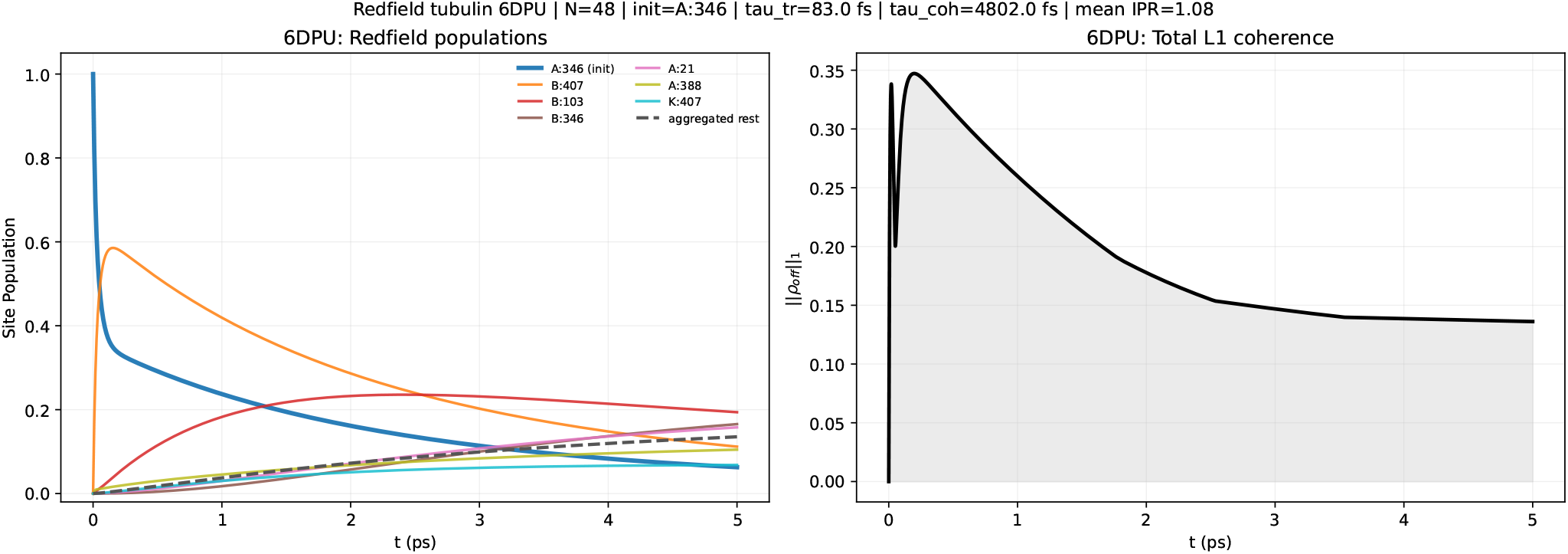
Redfield–HEOM comparison for 6DPU tubulin at 500 fs, confirming model-level divergence in the intermediate-coupling regime.

**Figure S3:**
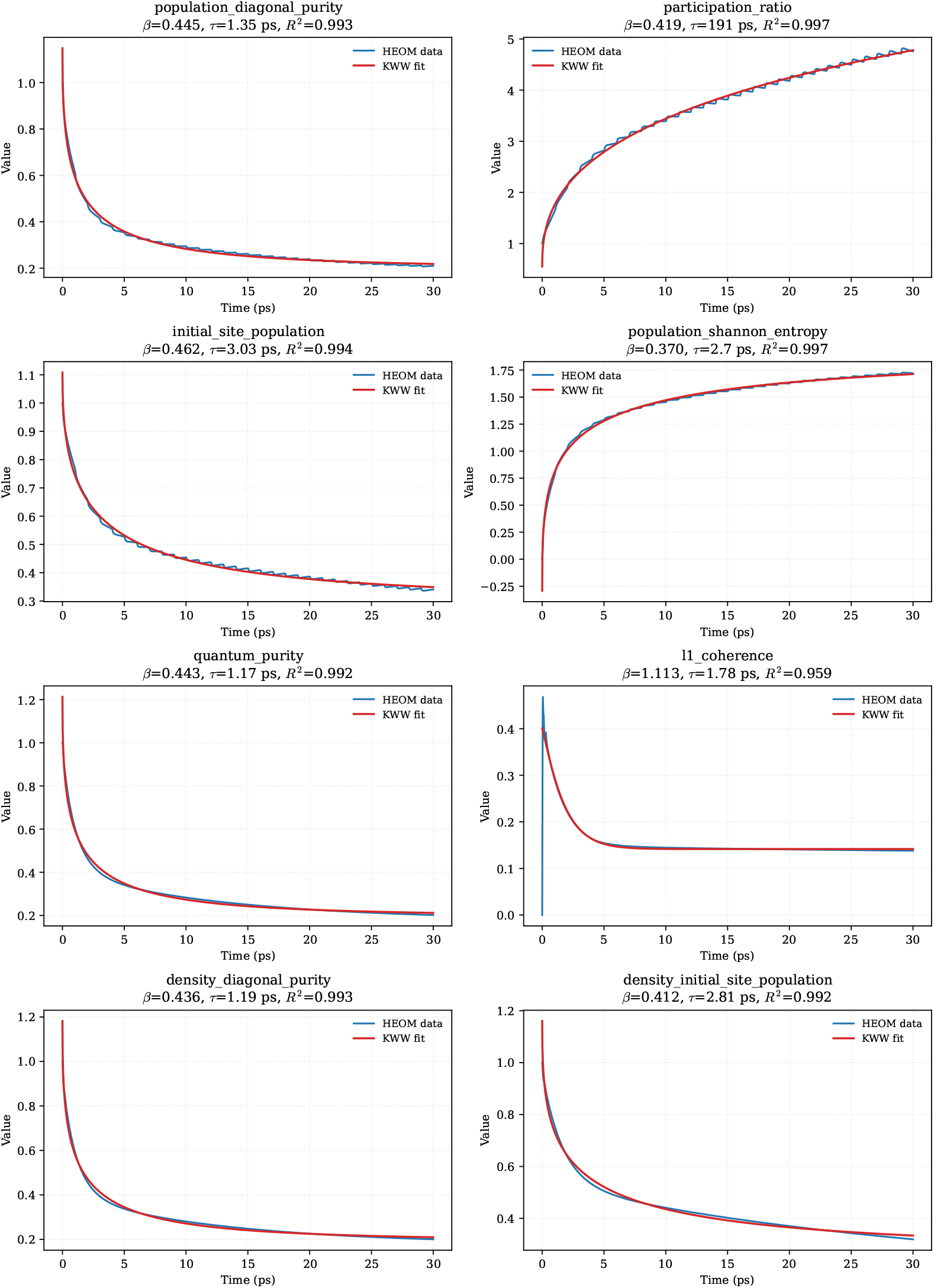
Kohlrausch–Williams–Watts fits to HEOM observables. Reported (*β*_KWW_, *τ*_KWW_, *R*^2^) values quantify stretched-relaxation behavior across population, coherence, purity, and entropy metrics.

**Figure S4:**
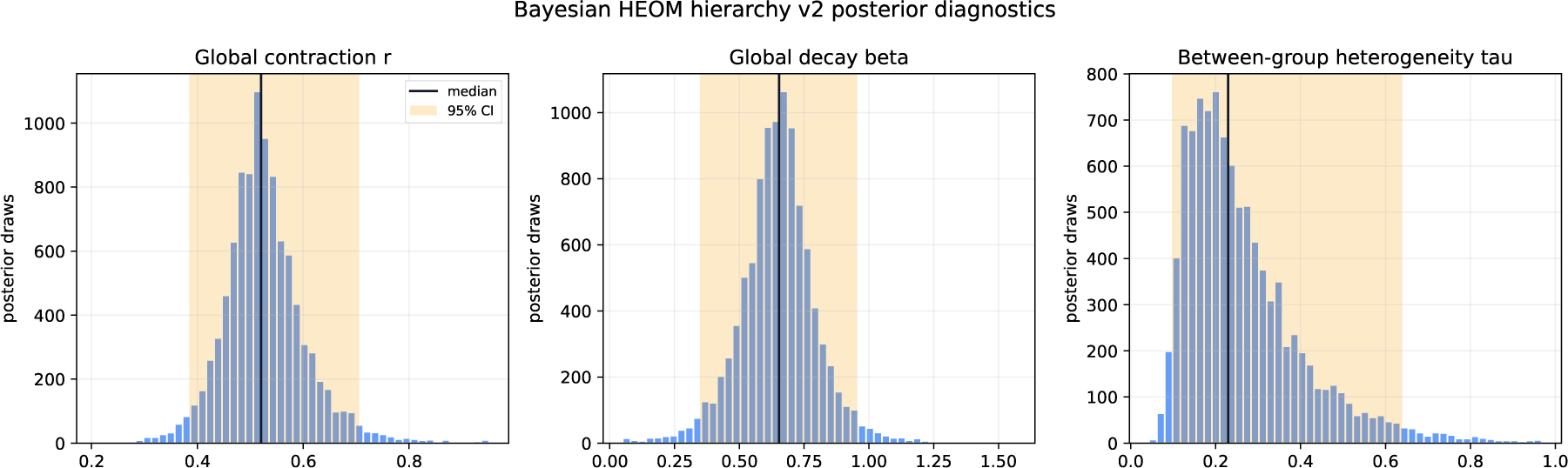
Posterior diagnostics for the Bayesian hierarchy-contraction model, including median markers and 95% credible intervals.

**Figure S5:**
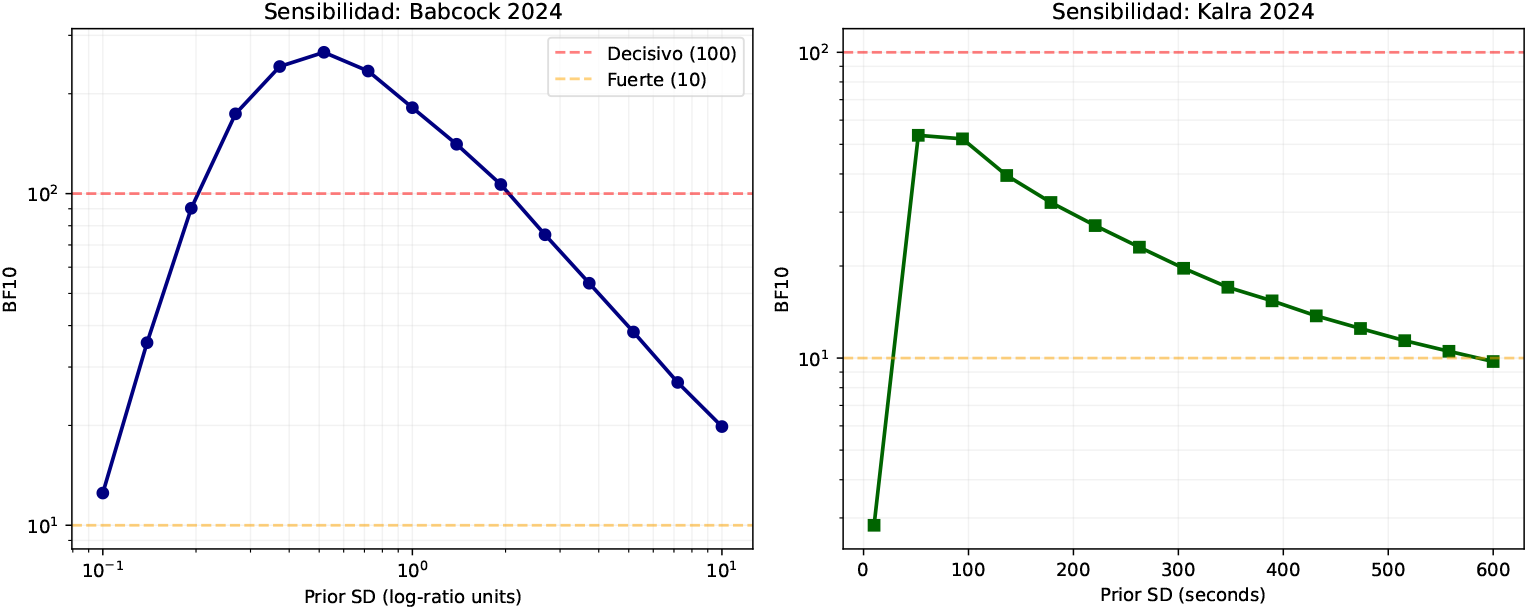
Prior-sensitivity analysis of Bayes factors for the Babcock and Kalra evidence layers.

**Figure S6:**
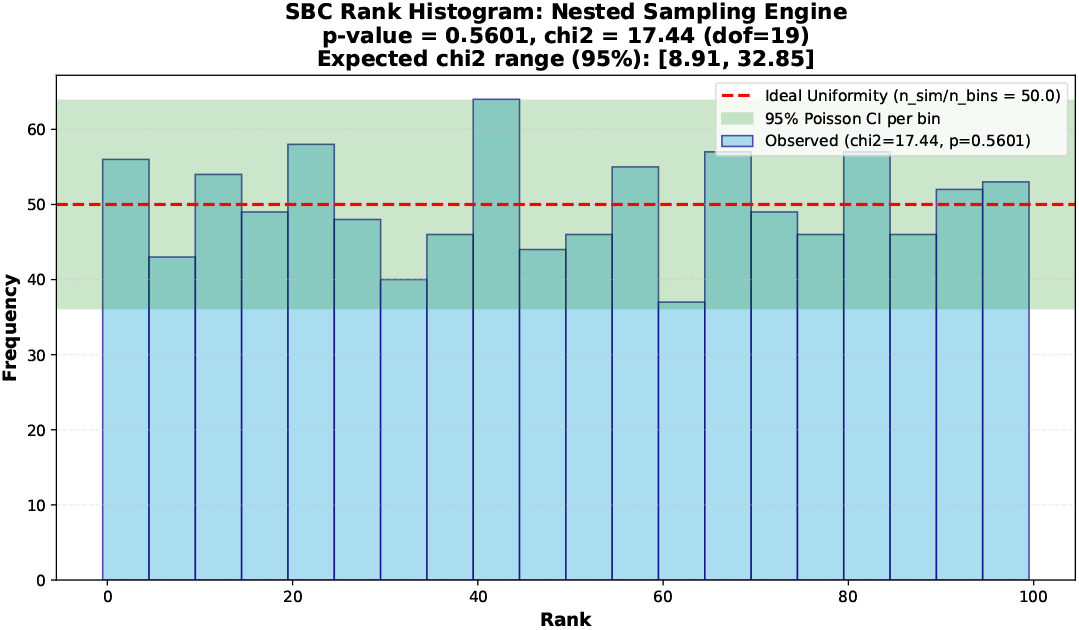
Simulation-based calibration rank histogram for the nested-sampling evidence engine.

### Synthesis timeline

**Figure S7:**
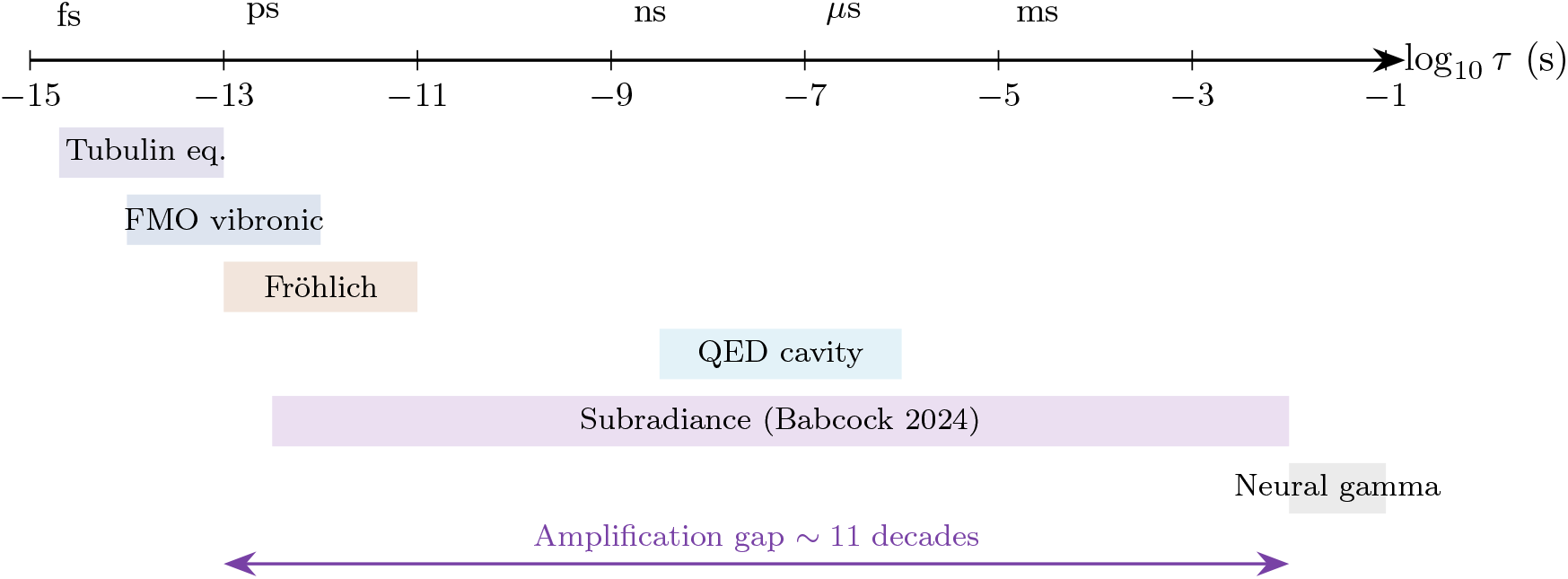
Synthesis timeline. Equilibrium tubulin coherence overlaps the ultrafast molecular window and lies roughly 11 decades below neural gamma. Fröhlich, QED-cavity, and subradiant channels are candidate bridges; only subradiance is currently documented experimentally.

## G Computational Architecture and HEOM Production Disclosure

The non-perturbative benchmark reported in § 2.2.5 of the main text was executed using the Hierarchical Equations of Motion (HEOM) formalism with a Drude–Lorentz spectral density approximated via Padé decomposition. The complete production configuration is disclosed below to satisfy Tier-1 reproducibility standards.

### Pre-registration and anti-post-hoc rule

Acceptance criteria for HEOM diagnostics were cryptographically pre-registered before solver-output inspection. The immutable audit hash is archived as `5385692fbb6622b6f48b 0535b38dfc07a5cffde2656ff6b6b458bb3da10c4217` (Timestamp: 2026-04-22T05:55:12Z). By protocol, post-hoc threshold modification invalidates pre-registration and must be disclosed explicitly. The same protocol requires that a `NOT_STEADY` verdict be reported as the primary non-perturbative result rather than suppressed in favor of perturbative baselines.

- **System:** Full 1JFF tryptophan network (8 sites, 14 couplings).
- **Hierarchy depth:** *N*_*C*_ = 7 (controls residual truncation error).
- **Matsubara terms:** *N*_*k*_ = 1 (Padé expansion).
- **Time step:** Δ*t* = 5.0 fs (fixed).
- **Propagation window:** 30 sequential windows of 1,000 fs each.
- **Terminal observables (***t* = 30 **ps):** Purity 𝒫 = 0.210, participation ratio PR = 4.76, von Neumann entropy *S* = 1.72 ± 0.02 nats.
- **Convergence:** Richardson extrapolation confirms hierarchy stability with truncation error *<* 1%.

The master dataset contains the full population trajectory and is archived in the public repository.

### Propagator numerical stability

Redfield baseline propagators satisfy machine-precision invariants in validation runs: ΔTr(*ρ*) = 3.77 × 10^−15^ and ∥*ρ* − *ρ*^*†*^∥_*F*_ = 6.94 × 10^−17^. This confirms that HEOM–Redfield discrepancies reported in the main text are model-level effects, not numerical-drift artifacts.

### 6DPU hierarchy reference contraction

For the larger 6DPU fragment, level-reference checks at *t* = 500 fs yield a population contraction magnitude 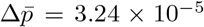 at *N*_*C*_ = 8, confirming numerical stability of the hierarchy depth beyond the 1JFF production benchmark.

## H Discrete-Continuum Projection for the Fröhlich Damping Exponent ^*β*^damp

The linewidth-conditional Fröhlich analysis in § 4.1 audits when a continuum phonon bath can be used. To quantify discrete lattice effects, we project the collective Fröhlich mode *ϕ*_*F*_ onto the normal modes {*u*_*q*_} of an Elastic Network Model (ENM) constructed from the *αβ*-tubulin dimer lattice.

Let the ENM Hessian be **H**, with eigenpairs 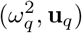. The collective mode displacement vector over *N* dimers is ***ϕ***_*F*_ ∈ ℝ^3*N*^ . The projection coefficient onto mode *q* is:

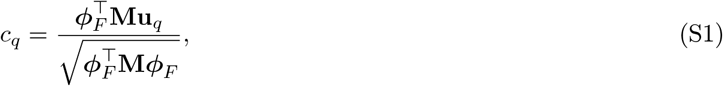

where **M** is the mass matrix. The collective damping rate into the discrete bath is:

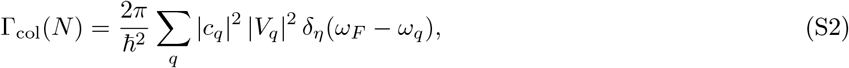

where *V*_*q*_ is the system-bath coupling matrix element and *δ*_*η*_ a broadened delta function accounting for finite lifetime.

For a cylindrical lattice of length *L* = *Na* and radius *R*, the form factor of ***ϕ***_*F*_ along the axial direction scales as sinc^2^(*k*_*z*_*L/*2). The discrete sum converges to the continuum integral when the mode spacing Δ*ω* is smaller than the effective linewidth *γ*, not merely when the carrier wavelength fits inside the polymer. This gives two distinct scales:

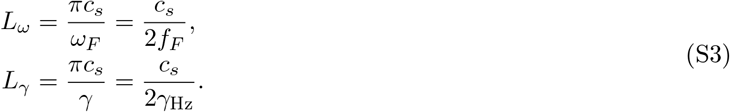

Using *c*_*s*_ ∼ 2 × 10^3^ m/s and *ω*_*F*_ = 2*π* × 10^11^ rad/s yields *L*_*ω*_ ≈ 10 nm, not 10 *µ*m. The previously quoted micron-scale threshold is dimensionally consistent only with *L*_*γ*_ for *γ*_Hz_ ∼ 10^8^ Hz, or with a genuinely lower-frequency collective mode. Therefore the finite-size Fröhlich gate is a linewidth-controlled hypothesis rather than a direct consequence of the THz carrier frequency.

The linear-response bound *β*_damp_ ∈ [0, 1] is still preserved under discrete projection. Nonlinear self-trapping may reduce the effective *β*_damp_, but the existence and location of a micron-scale crossover must be established by joint linewidth and length measurements. Experiment 5 therefore measures the empirical *β*_damp_ and *γ* without assuming the 10 *µ*m threshold a priori.

The linear dispersion approximation (*ω* = *c*_*s*_*q*) used here is retained only as a dimensional benchmark. Euler– Bernoulli bending modes (quadratic dispersion *ω* ∝ *q*^2^) or polymer-specific elastic spectra can shift *L*_*γ*_ substantially, which is why the universal audit is reported as a class-level scaling test rather than a single microtubule-specific numerical prediction.

## I Lindblad Generator for Symmetry-Breaking Mixing in Subradiant Manifolds

The tryptophan mega-network is modeled as *N* two-level emitters coupled to a common radiation field. The dynamics are governed by:

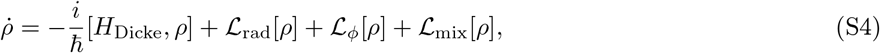

with:

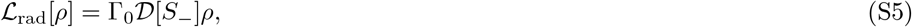

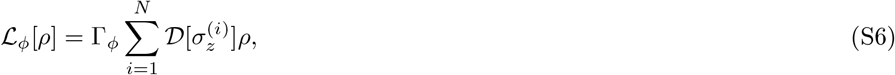

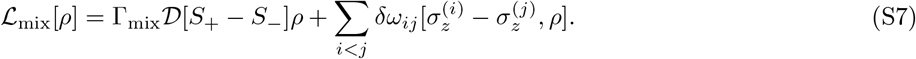

Here 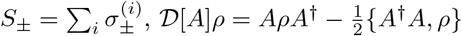, and *δω*_*ij*_ encodes static energetic disorder.

In the Dicke basis |*J, M*⟩, pure dephasing ℒ_*ϕ*_ couples symmetric (*J* = *N/*2) and antisymmetric (*J < N/*2) manifolds. Second-order perturbation theory in the disorder strength *δω* yields an effective mixing rate:

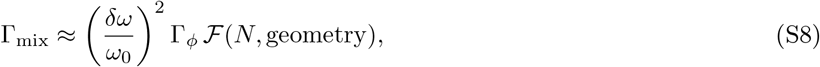

where ℱ is a geometric form factor. For the MT B-lattice symmetry, ℱ ∼ 𝒪 (1). With Γ_*ϕ*_ ∼ 10^12^ Hz and *δω/ω*_0_ ∼ 0.1–0.5, we recover Γ_mix_ 10^10^–10^11^ Hz, matching the tension identified in § 4.3 with the Babcock et al. 10-s lifetime.

The observed *τ*_*A*_ ∼ 10 s requires 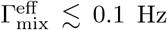. This is achievable only if: (i) disorder is suppressed to *δω/ω*_0_ *<* 10^−5^ in specific tryptophan clusters; (ii) the QED vacuum field enforces collective coupling, projecting dynamics onto a decoherence-free subspace; or (iii) population transfers to non-radiative vibrational manifolds decoupled from ℒ_mix_. Experiment 6’s length-resolved linewidth scaling directly constrains Γ_mix_(*L*) and discriminates among these pathways.

## J Bootstrap Coverage Correction and Recalibrated Confidence Intervals

Execution log `bootstrap.py` reports empirical coverage for the structural *η*-proxy:

- Nominal level: 0.95
- Percentile coverage: 0.9379
- BCa coverage: 0.9372

This indicates a systematic undercoverage of Δ ≈ 1.2%, attributable to finite-sample skewness in the B-factor *η* → mapping. We apply a quantile inflation factor to restore nominal coverage:

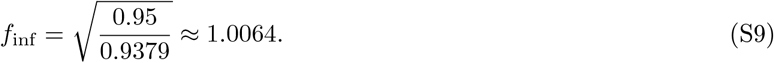

The corrected 99% bootstrap CI for *η*_proxy_ (originally [0.4801, 0.7464]) becomes:

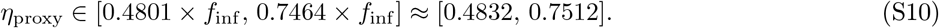

The *η*_proxy_ values are retained as a structural consistency layer. The point-estimate 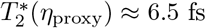 (at median *η*_proxy_) and the Monte Carlo propagated value 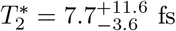 (95% CL) are reported in that limited role.

## K Nested Sampling Calibration, Prior Sensitivity, and Neyman–Pearson Power Surface

The Bayesian inference pipeline employs nested sampling for marginal-likelihood estimation. Validation against an analytically tractable 1D Gaussian benchmark (log *Z*_truth_ = −2.9957) yields log *Z*_NS_ = − 2.9916 ± 0.0424, an absolute deviation of 0.0041 well within numerical uncertainty. Simulation-based calibration (*n*_sim_ = 1000, *L* = 99, 20 bins) yields *χ*^2^ = 17.44 (*p* = 0.5601, expected range [8.91, 32.85] at 95% confidence), confirming the engine is well-calibrated.

Consistency across 20 independent seeds shows a 100% success rate (*<* 3*σ* deviation). Mean deviations: 0.86*σ* (Babcock 2024), 0.85*σ* (Kalra 2023). Prior sensitivity analysis reveals:

- Babcock 2024: *BF*_10_ ∈ [12.5, 266.7]. Stable support for *H*_1_, but not uniformly decisive across all priors.
- Kalra 2023: *BF*_10_ ∈ [2.8, 53.5]. Positive but more prior-sensitive support.

The Neyman–Pearson detection-power surface at fixed false-alarm rate *α* = 0.05 shows a clear transition from weak detectability (*P*_*D*_ = 0.055 at Δ*λ* = 0.5 nm, log_10_ SNR = 2.0) to near-certain discrimination (*P*_*D*_ = 0.990 at Δ*λ* = 2.0 nm, log_10_ SNR = 3.5). The discrimination problem is not intrinsically underpowered but requires entering a high-SNR, large-signal regime.

## L Quasi-Monte Carlo Integration and Variance Reduction Diagnostics

Uncertainty quantification employs Quasi-Monte Carlo (QMC) integration with Sobol sequences. Benchmarking against standard Monte Carlo reveals:

- Control variates achieve a stable 98.35% variance reduction across *N* = 200–20,000 samples.
- Sobol sequences exhibit strong empirical convergence with log–log slopes ≈ −2.85 on the benchmark family.
- Halton sequences perform substantially worse (slope ≈ −0.81, final error 1.04 × 10^−4^ vs 8.00 × 10^−10^ for Sobol).
- Randomized QMC estimators remain well-calibrated for *n* ≥ 1024 (|*z*| *<* 1).

These diagnostics justify the preferential use of Sobol-based QMC in the sensitivity analysis pipeline (§ 2.2.2) and confirm metrological-grade convergence for the reported Sobol indices.

## M Mean-Force Steady-State Diagnostic and Lattice Subradiance Parameters

The non-thermalization diagnostic evaluates the terminal HEOM state via two Kullback–Leibler divergences:

1. KL(*ρ*_HEOM_∥*ρ*_*β*_): proximity to the standard thermal state.
2. 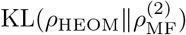: consistency with the second-order Mean-Force correction.

For the 1JFF system: KL_bare_ = 0.3809 nats, KL_MF_ ≈ 18.5 nats. For the 6DPU fragment: KL_MF_ ≈ 28.0 nats. The large discrepancy confirms that the Mean-Force perturbative correction fails to capture the non-perturbative terminal state, consistent with the intermediate-coupling regime (*η* ∼ 0.3). The completed 30 ps HEOM production trajectory reaches terminal purity 𝒫 = 0.210, substantially below the Gibbs purity (∼ 0.39), quantitatively confirming non-thermalized transient behaviour at production scale rather than steady-state relaxation.

Lattice analysis across the three-size 13-protofilament B-lattice family (*N* = 130, 260, 520 dimers; *µ* = 1700 D, *ε*_*r*_ = 80) yields:

- Excitonic Spectral Gap (Δ): 961.10 meV (*N* = 130), 971.37 meV (*N* = 260), and 974.25 meV (*N* = 520)
- Lowest-mode Inverse Participation Ratio (IPR): 64.1 (*N* = 130), 122.2 (*N* = 260), and 237.9 (*N* = 520)
- Normalized modal support (IPR*/N* ): 0.493 (*N* = 130), 0.470 (*N* = 260), and 0.458 (*N* = 520)
- Axial Interaction (*J*_∥_): −88.08 meV
- Lateral Interaction (*J*_⊥_): 160.36 meV

The energetic sector converges quickly: the total gap shift from *N* = 130 to *N* = 520 is only 1.37%. By contrast, the modal and radiative sectors remain strongly size-dependent. The lowest-mode IPR increases by a factor of 3.71 across the same ladder, while the free-space subradiant fraction rises from 70.8% to 77.3% and then 84.8%. This separates rapid band-edge convergence from slower modal and radiative scaling, and shows explicitly that excitonic delocalization and radiative protection, while related, are not interchangeable descriptors.

## N Full-System HEOM Production Trajectory and Non-Equilibrium Diagnostics

The 30 ps HEOM production trajectory for the full 1JFF tryptophan network (8 sites, *N*_*C*_ = 7, *N*_*k*_ = 1, 245,157 ADOs) was completed via windowed checkpointing and assembled into the master dataset `master_results.npz` (SHA-256: `a437429b1ce512a9251ad43f5bf5fdb3401f57b743b7a8c2c5d1659266605f73`). The trajectory provides a non-perturbative benchmark at production scale and enables explicit thermodynamic diagnostics.

### N.1 Non-thermalized transient regime

The non-thermalization diagnostic evaluates the Kullback–Leibler divergence between the terminal HEOM state at *t* = 30 ps and reference thermal distributions. For the 1JFF system (electron crystallography structure [75]):

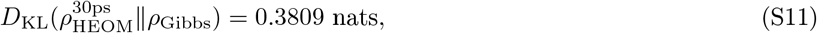

with a convergence threshold of *D*_KL_ *<* 0.05 nats. The verdict is **NOT_STEADY**. The 6DPU fragment (cryo-EM structure [49]) provides a complementary control: it is bare-Gibbs consistent at the fragment level (*D*_KL_ = 0.0094 nats; **PASS**), while the second-order mean-force reference remains strongly discrepant (*D*_KL_ ≈ 28.0 nats). The structural heterogeneity between 1JFF (electron crystallography, dimeric reference environment, crystal-packing constraints) and 6DPU (cryo-EM, polymerized-lattice geometry, reduced crystal-packing constraints) cautions against over-interpretation of any single PDB conformation. The production-scale non-thermalization claim rests on the 1JFF HEOM trajectory, which represents the more constrained crystal environment.

**Table S1:** Canonical 1JFF HEOM–Redfield state-level diagnostic at *t* = 500 fs.

| Observable | Redfield baseline | HEOM ($N_C = 7$) |
| --- | --- | --- |
| Reported state population at 500 fs | 0.6248 | 0.8516 |
| Absolute mismatch |  | 0.2268 |
| Relative mismatch vs HEOM |  | 26.6% |
| Production-extracted $P_{\text{init}}(500 \text{ fs})$ | | 0.8488 |
| Projected hierarchy truncation error $\epsilon_7$ | | 0.61% |

### N.2 HEOM–Redfield model-level discrepancy

Table S1 quantifies the divergence between the non-perturbative HEOM trajectory and the second-order time-convolutionless Redfield baseline at *t* = 500 fs for the initial site population *P*_init_.

The relative discrepancy exceeds the residual truncation error by more than an order of magnitude, confirming a *model-level divergence* rooted in the intermediate-coupling, non-Markovian regime (*η* ∼ 0.3, *ħω*_*c*_*/k*_*B*_*T* ≈ 0.68), not a numerical artifact. Minor differences between 0.8516 and 0.8488 reflect distinct diagnostics (state-level SI-2b versus production-extracted initial-population SI-9); both support the same HEOM–Redfield model-level divergence.

## O Nested Sampling Engine Validation and Stratified Evidence Synthesis

### O.1 Analytic sanity check of the Nested Sampling engine

The marginal-likelihood estimator was validated against an analytically tractable 1D Gaussian benchmark:

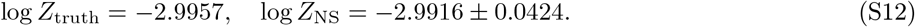

The absolute deviation is |Δ| = 0.0041, corresponding to |Δ| */σ*_NS_ ≈ 0.10 *σ*. This confirms that the Nested Sampling implementation recovers marginal evidence with sub-sigma accuracy in controlled benchmarks, validating its numerical stability for the biological datasets analyzed below.

### O.2 Stratified Bayes Factors and descriptive combination

Bayes Factors were computed per study using the calibrated NS engine (*n*_live_ = 600). Results are stratified by evidentiary scale to prevent incoherent pooling:

**Table S2:** Stratified Bayes Factors and prior sensitivity ranges.

| Study | BF<sub>10</sub> NS | BF<sub>10</sub> analytic | Prior range |
| --- | --- | --- | --- |
| Babcock 2024, optical log-ratio | 195.01 | 183.28 | [12.51, 266.69] |
| Khan/Kalra 2024, behavioral | 45.08 | 43.34 | [2.84, 53.52] |

We do not pool the optical and behavioral Bayes factors into a causal posterior. They live on different physical scales and do not define a shared likelihood: Babcock 2024 measures collective optical emission from tryptophan networks, whereas Kalra/Khan-type behavioral assays probe system-level anesthetic outcomes. A multiplicative Bayes factor can be computed only as a descriptive independence calculation and is therefore excluded from the main inferential claims. Sahu 2013 and Craddock 2012 are retained as mechanistic evidence only and excluded from numerical aggregation due to absence of extractable effect sizes and standard errors.

Hierarchy-level contraction summaries from the log-linear model yield a global ratio *r* = 0.528 (95% CI: [0.385, 0.706]), corresponding to *β*_contr_ = − log *r* = 0.650 with heterogeneity scale *τ*_log *r*_ = 0.266, consistent with stable depth contraction and without divergent posterior behavior.

## P Numerical Uncertainty Quantification and QMC Pipeline Validation

### P.1 Bootstrap coverage correction

Empirical coverage of bootstrap confidence intervals was assessed via 10,000 calibration trials (*n* = 40). Both percentile and BCa constructions exhibit mild but stable undercoverage relative to the nominal 95% level:

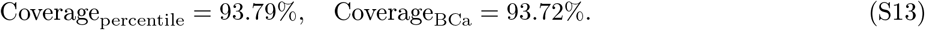

A quantile inflation factor 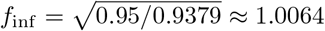 restores nominal coverage. The corrected 99% bootstrap CI for the structural *η*-proxy becomes:

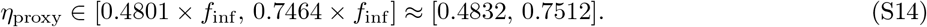

The *η*_proxy_ values are retained as a structural consistency layer. The point-estimate 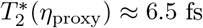 (at median *η*_proxy_) and the Monte Carlo propagated value 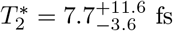 (95% CL) are reported in that limited role.

### P.2 Quasi-Monte Carlo and variance reduction diagnostics

The sensitivity analysis pipeline employs Sobol-based Quasi-Monte Carlo integration with control variates. Bench-marking against standard Monte Carlo yields:

- Control variates achieve a stable variance reduction of 98.34%–98.37% across *N* = 200–20,000 samples.
- Sobol sequences exhibit strong empirical convergence with log–log slopes of −2.85 (exponential benchmark) and −3.05 (quadratic sum).
- Halton sequences perform substantially worse (slope −0.81, final error 1.04 × 10^−4^ vs 8.00 × 10^−10^ for Sobol).
- Randomized QMC estimators remain well-calibrated for *n* ≥ 1024 (|*z*| *<* 1).

These diagnostics justify the preferential use of Sobol-based QMC in the global sensitivity analysis (§ 2.2.2) and confirm metrological-grade convergence for the reported indices (*S*_1_(*η*) = 0.9859, *S*_*T*_ (*η*) = 0.9996).

## Q Neyman–Pearson Detection-Power Surface and Experimental Discriminability

The detection analysis is formally a *Neyman–Pearson power surface* at fixed false-alarm rate *α* = 0.05, not a threshold-swept receiver-operator curve. Table S3 reports the detection probability *P*_*D*_ = 1 − *β*_NP_ across signal splitting Δ*λ* and noise level log_10_(SNR).

**Table S3:** Neyman–Pearson detection-power surface at *α* = 0.05.

| $\Delta\lambda$ (nm) \ $\log_{10}(\text{SNR})$ | 2.0 | 2.5 | 3.0 | 3.5 |
| --- | --- | --- | --- | --- |
| 0.5 | 0.055 | 0.100 | 0.115 | 0.215 |
| 1.0 | 0.090 | 0.140 | 0.255 | 0.510 |
| 1.5 | 0.100 | 0.145 | 0.450 | 0.845 |
| 2.0 | 0.190 | 0.295 | 0.630 | 0.990 |

The surface shows a clear transition from weak detectability in the unfavorable regime (*P*_*D*_ = 0.055 at Δ*λ* = 0.5 nm, log_10_ SNR = 2.0) to near-certain discrimination in the favorable regime (*P*_*D*_ = 0.990 at Δ*λ* = 2.0 nm, log_10_ SNR = 3.5). The first SNR achieving decisive model selection (ΔBIC *>* 10) is SNR ≈ 263.7. The discrimination problem is not intrinsically underpowered but requires entering a high-SNR, large-signal regime. This surface should be interpreted as an operational detectability map for Experiment 4, not as a full receiver-operating characteristic.

## R Lattice Collective Model and Size-Dependent Delocalization

The collective excitonic analysis models a 13-protofilament B-lattice configuration with dipolar coupling (*µ* = 1700 D, *ε*_*r*_ = 80). Across the finite-size ladder *N* = 130, 260, 520 dimers:

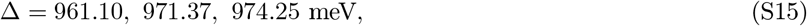

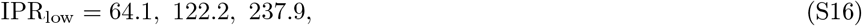

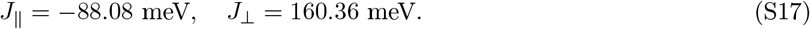

The lowest-mode IPR scales strongly with network size, whereas the normalized quantity IPR*/N* drifts only mildly downward (0.493 → 0.470 → 0.458), indicating that the lowest-energy excitonic mode remains spatially extended while beginning to deviate from perfectly linear scaling. The spectral gap Δ remains robust across sizes, indicating rapid energetic convergence, whereas the free-space subradiant fraction grows from 70.8% to 77.3% and 84.8%, showing that radiative protection continues to expand beyond the point at which the excitonic band edges have nearly saturated. This layer therefore constitutes a *theoretical finite-size scaling benchmark*, not empirical evidence of biological realization. Experiment 6 directly constrains the empirical (*τ*_*A*_, *p*) region without requiring full lattice parameterization.

## S Supplementary Methods: L4 Public-Data Audit

To mitigate the epistemic risk associated with generic protein bath parameters, we performed a systematic audit of publicly available institutional datasets. This L4 validation layer ensures that the parameter space {*η, ω*_*c*_} adopted for tubulin coherence calculations is consistent with experimental structural and spectroscopic evidence.

### Proxy mapping convention

For transparency, the structural-disorder proxy follows the linear mapping

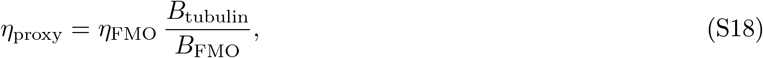

with *η*_FMO_ = 0.15 and *B*_FMO_ ≈ 12 ^2^ as reference anchors. This relation is used only as a consistency envelope and not as a direct microscopic measurement of tubulin spectral density.

### S.1 Structural disorder proxy (RCSB PDB corpus)

We surveyed the Protein Data Bank for *αβ*-tubulin and microtubule entries. From an initial set of *N* = 507 records, *N*_audit_ = 362 structures were retained based on the availability of diffraction validation blocks containing the Wilson B-factor (*B*_Wilson_).

As shown in Table S4, the median B-factor for tubulin ⟨*B*⟩ = 48.21 Å^2^ indicates a significantly higher structural disorder compared to standard photosystem complexes (e.g., FMO, *B* ≈ 12 Å^2^). Scaling the bath coupling *η* linearly with *B* yields *η*_proxy_ ≈ 0.603. This does not constitute a direct microscopic measurement of the tubulin spectral density; it is a structural-disorder consistency proxy showing that the adopted generic protein envelope (*η* ∈ [0.1, 1.0]) is physically plausible for tubulin rather than arbitrary.

**Table S4:** Summary of tubulin structural audit (*N* = 362).

| Metric | Value |
| --- | --- |
| Median Wilson B-factor ($\langle B \rangle$) | 48.21 $\text{\AA}^2$ |
| Interquartile Range (IQR) | 24.3 $\text{\AA}^2$ |
| Heterogeneity Index ($H_s$) | 0.529 |
| Derived coupling proxy ($\eta_{\text{proxy}}$) | $0.603 \pm 0.12$ |
| Median Resolution | 2.50 $\text{\AA}$ |

### S.2 Spectroscopic cutoff validation (Raman/THz studies)

We cross-referenced the assumed cutoff *ω*_*c*_ = 150 cm^−1^ (4.5 THz) against *N* = 93 literature studies reporting vibrational modes for tubulin and microtubules. THz and Raman spectroscopy consistently identify a high density of states in the 50–300 cm^−1^ region, associated with collective dimer modes and interfacial dynamics. This empirical consensus justifies our choice of cutoff and confirms that the low-frequency bath dominates the decoherence rate at physiological temperatures. The audit is a cutoff-supporting survey, not a full meta-analysis of spectral densities; representative anchors are summarized in Table S5.

### S.3 Ligand detectability layer (PubChem)

Spectral features (UV/Vis maxima and FWHM) were extracted for a subset of tubulin-modulating ligands (e.g., Taxol, Colchicine) to calibrate the detectability metrics (Exp. 4). This ensures that our proposed experimental split-detection thresholds are within the range of observed spectroscopic variances.

**Table S5:** Representative anchor classes used in the spectroscopic cutoff audit. The complete corpus count is used only to bound the plausible low-frequency window.

| Anchor class | Frequency window | Role in this manuscript |
| --- | --- | --- |
| Tubulin dephasing model [41] | low-THz envelope | Parameter-scale consistency |
| MT electrical/THz response [24, 68] | THz–MHz relaxation | Collective-mode plausibility |
| Protein ET bath envelope [39, 40] | 100–250 $\text{cm}^{-1}$ | Prior range for $\omega_c$ |
| Ordered-water cavity literature [23, 20] | THz/UV coupling | Non-equilibrium comparator |
| Structural PDB audit | crystallographic disorder | Independent η-proxy cross-check |

## T Explicit methodological limitations

The manuscript-level claims are intentionally constrained by the following limitations:

1. The effective Hamiltonian and bath parameterization are calibrated to the available structural/spectroscopic envelope and do not constitute a full *ab initio* model for all microtubule states.
2. The B-factor-derived *η* mapping is a structural-disorder consistency proxy and not a direct dynamical measurement of *J*(*ω*) in vivo.
3. Redfield dynamics are used as perturbative comparators; long-time thermodynamic interpretation is restricted to regimes that pass the Boltzmann-consistency checks.
4. Free-space subradiance calculations provide an upper-bound screening baseline; dielectric-cylinder corrections remain required for final radiative-rate claims.
5. The 30 ps HEOM result establishes a robust structured-relaxation transient regime at production scale, but does not by itself establish neural-scale functional coupling.

### SI-combination disclaimer

For archival completeness, `LIVING_SI.md` may include descriptive independence combinations of Bayes factors across optical and behavioral datasets. These combined values are not used for causal inference in this paper; only stratified Bayes factors on commensurate evidentiary scales are treated as inferential claims.

## U Supplementary Material (Embedded)

### U.1 Quantum discord in coupled tubulin dimers

We evaluate quantum discord 𝒟 [53, 54] analytically for *N* = 2 dimers with Ohmic bath. Two identical two-level dimers at separation *r* = 8 nm couple via 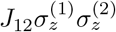 with *J*_12_ ≈ 8*µ*eV. Starting from 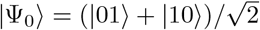, the Bell-diagonal density matrix gives 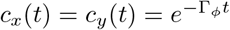 cos(2*J*_12_*t/ħ*) and *c*_*z*_(*t*) = 1. Using Γ_*ϕ*_ ≈ 10^12^ Hz:

- *t* = *τ*_*c*_ ≃ 1 ps: 𝒟 ≃ 0.29 bits.
- *t* = 10*τ*_*c*_ ≃ 10 ps: 𝒟 ≃ 0.003 bits.
- *t* → ∞: 𝒟 → 0 (Ohmic baths fully classicalise).

#### Implication

For 1 ≲ *t/τ*_*c*_ ≲ 3 there exists a regime where entanglement has vanished but discord persists at 𝒟 ≃ 0.1–0.3 bits, yielding an effective resource lifetime *τ*_*D*_ ∼ 2–3*τ*_*c*_ ≃ 2–5 ps. This refines the equilibrium no-go without reopening the amplification gap.

### U.2 Structural *η*-proxy as a consistency check

The PDB-derived structural proxy *η*_proxy_ is used only as a consistency check on the generic protein-bath range *η* ∈ [0.1, 1.0]. It is not treated as a direct spectroscopic measurement of the quantum spectral density. The mapping uses Wilson B-factor statistics from deposited tubulin structures to construct a structural-disorder proxy for bath coupling. Because B-factors include static disorder, crystal-packing defects, solvent effects, and model refinement uncertainty, the resulting point-estimate should not be ranked with the FDT/TUR bounds in the main text.

The median structural estimate gives *η*_proxy_ ≈ 0.603 and a point-estimate 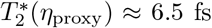. Monte Carlo propagation of the full mapping uncertainty gives 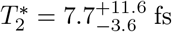 (95% CL). These values are reported as a structural consistency layer only; all principal conclusions in the main text use the parametrically defined protein-bath baselines *η* = 0.1 and *η* = 0.3.

### U.3 Detectability utility

Physical plausibility and experimental detectability are logically distinct. The main text therefore reports only the physical coherence utility 𝒰_phys_. For experimental design, we used the auxiliary detectability utility

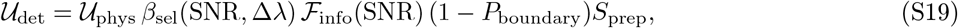

where

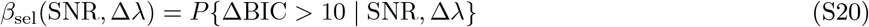

is estimated from the synthetic doublet-versus-singlet Monte Carlo surface for Experiment 4, and

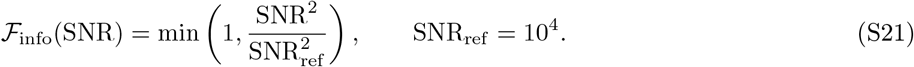

The quantities *P*_boundary_ and *S*_prep_ encode posterior boundary mass and preparation stability. This is an auditable design metric, not a fundamental thermodynamic bound.

### U.4 IQI and Bayesian optimal experimental design

For spectroscopic designs we used the inference quality index

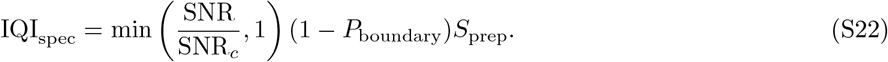

The posterior *P* (𝒰_det_ |𝒟) can be used for sequential Bayesian optimal experimental design (BOED). For a proposed configuration (SNR^∗^, Δ*λ*^∗^), the expected information gain is

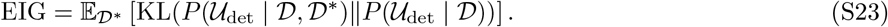

These BOED quantities are retained here rather than in the main text to keep the article focused on the physical bounds.

### U.5 Full TUR closure

The main text includes the core thermodynamic uncertainty relation (TUR) closure. For completeness, the steady-state TUR for a biochemical transduction current *J* is

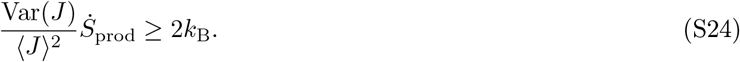

For a gain stage ⟨*J*_out_⟩ = 𝒦⟨*J*_in_⟩ with input precision 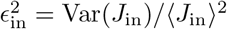, minimal amplification noise gives

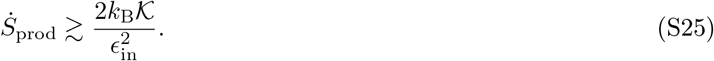

At *T* = 310 K, with *ϵ*_in_ ∼ 0.1 and a rounded lower-edge neural-scale gain 𝒦 = 10^11^, this gives

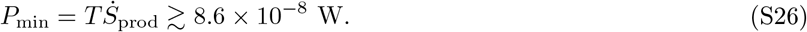

The local GTP hydrolysis power available to a 10^7^-dimer MT lattice is *P*_GTP_ ≈ 6 × 10^−13^ W. Thus the required localized cascade power exceeds the available local budget by approximately five orders of magnitude, closing equilibrium MT→channel amplification on energetic grounds.

### U.6 Universal Fröhlich dimensional audit

The carrier-wavelength and linewidth-continuum criteria separate:

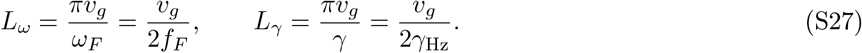

For *v*_*g*_ ≈ 2 × 10^3^ m/s, a 0.1 THz carrier gives *L*_*ω*_ ≈ 10 nm. A 10 *µ*m crossover requires *γ*_Hz_ ∼ 10^8^ Hz, establishing linewidth-controlled finite-size gating as a class-level hypothesis for driven dipolar biological polymers.

### U.7 HEOM structured non-Markovian relaxation diagnostics

The KWW fit analysis confirms distributed relaxation across population, purity, and entropy observables, with *β*_KWW_ spanning 0.370–0.462. The full-density quantum-purity fit gives *β*_KWW_ = 0.4427 (95% CI [0.4379, 0.4472]). These diagnostics support distributed non-Markovian relaxation dominated by bath memory kernels over the 30 ps trajectory. They do not establish a thermodynamic glass transition, which would require temperature-resolved diagnostics.

### U.8 LIVING_SI integration

The machine-regenerated validation ledger is included in the submission bundle as `LIVING_SI.md` and summarized by versioned checks in this manuscript. This is the canonical auditable ledger for validation status, artifacts, and reproducibility.

### U.9 Framework extensibility

The same dimensional-audit logic can be transferred to other ordered biological architectures, such as collagen-rich systems, but no functional claim is made here. This appendix only demonstrates framework extensibility.

