## Supplementary material for "Beyond Redfield: Thermodynamic Bounds and Non-Perturbative Quantum Dynamics in Tubulin Networks": Interactive Epistemic Graph: epistemic_graph.html

Interactive Epistemic Map of Claims, Evidence, and Falsification


### Interactive Epistemic Map of Claims, Evidence, and Falsification

Claim → evidence → validation ledger → falsifier → scope boundary

☰ Filters
♟ Inspector


Validation ledger

Scale separation

Visible nodes / edges

Target timescale


##### Lenses

##### Quick views

📉 Failed routes
🔬 Surviving candidates
🧪 Falsifiers
🚧 Scope boundaries
✅ Ledger‑backed
🌐 All

Select a view to see relevant manuscript sections.

##### Data Clutter

Hide isolated nodes (validation checks)

##### Legend

Fails current constraints

Conditional / utility‑limited

Falsifiable candidate

Ledger‑supported

Scope boundary

+
−
⊡

#### Select a node

Click any node or edge to inspect its epistemic role.

##### Artifacts

None selected.

##### Metrics

None selected.
